# Calmodulin binding is required for calcium mediated TRPA1 desensitization

**DOI:** 10.1101/2024.12.11.627969

**Authors:** Justin H. Sanders, Kehinde M. Taiwo, Glory A. Adekanye, Avnika Bali, Yuekang Zhang, Candice E. Paulsen

## Abstract

Calcium (Ca^2+^) ions affect nearly all aspects of biology. Excessive Ca^2+^ entry is cytotoxic and Ca^2+^- mobilizing receptors have evolved diverse mechanisms for tight regulation that often include Calmodulin (CaM). TRPA1, an essential Ca^2+^-permeable ion channel involved in pain signaling and inflammation, exhibits complex Ca^2+^ regulation with initial channel potentiation followed by rapid desensitization. The molecular mechanisms of TRPA1 Ca^2+^ regulation and whether CaM plays a role remain elusive. We find that TRPA1 binds CaM best at basal Ca^2+^ concentration, that they co-localize in resting cells, and that CaM suppresses TRPA1 activity. Combining biochemical, biophysical, modeling, NMR spectroscopy, and functional approaches, we identify an evolutionarily conserved, high-affinity CaM binding element in the distal TRPA1 C-terminus (DCTCaMBE). Genetic or biochemical perturbation of Ca^2+^/CaM binding to the TRPA1 DCTCaMBE yields hyperactive channels that exhibit drastic slowing of desensitization with no effect on potentiation. Ca^2+^/CaM TRPA1 regulation does not require the N-lobe, raising the possibility that CaM is not the Ca^2+^ sensor, *per se*. Higher extracellular Ca^2+^ can partially rescue slowed desensitization suggesting Ca^2+^/CaM binding to the TRPA1 DCTCaMBE primes an intrinsic TRPA1 Ca^2+^ binding site that, upon binding Ca^2+^, triggers rapid desensitization. Collectively, our results identify a critical regulatory element in an unstructured TRPA1 region highlighting the importance of these domains, they reveal Ca^2+^/CaM is an essential TRPA1 auxiliary subunit required for rapid desensitization that establishes proper channel function with implications for all future TRPA1 work, and they uncover a mechanism for receptor regulation by Ca^2+^/CaM that expands the scope of CaM biology.

## INTRODUCTION

Calcium (Ca^2+^) is a unique ion since it regulates cell excitability and serves as a second messenger in signal transduction cascades involved in almost all aspects of cellular life^1^. Excessive Ca^2+^ entry is cytotoxic, and cells dedicate substantial resources to maintain a 20,000-fold lower cytoplasmic concentration than extracellular or endoplasmic reticulum stores^1^. Extracellular Ca^2+^ influx is facilitated by Ca^2+^-permeable ion channels whose activity must be tightly controlled to prevent spurious initiation of Ca^2+^ signaling pathways and apoptosis. In the peptidergic C fiber subset of peripheral sensory neurons, extracellular Ca^2+^ influx triggers pain signals and the exocytotic release of neuropeptides that initiate neurogenic inflammation and neuronal hypersensitivity^2–6^. These processes are believed to play a role in the transition from acute to chronic pain^7–9^. Identifying the Ca^2+^-permeable ion channels involved in initiating and maintaining neurogenic inflammation and determining their regulatory mechanisms could provide the basis for rational drug design to alleviate these symptoms.

A direct role for the wasabi receptor, TRPA1 (<u>T</u>ransient <u>R</u>eceptor <u>P</u>otential <u>A</u>nkyrin subtype <u>1</u>), in human pain has been illustrated by the discovery and characterization of genetic variants that are associated with painful disorders^10–12^. TRPA1 is a homotetrameric Ca^2+^-permeable non-selective cation channel that is expressed at the plasma membrane in a subset of peripheral sensory neurons originating from dorsal root, trigeminal, and nodose ganglia as well as non-neuronal tissues including airway epithelia, enterochromaffin cells, and cardiac tissue (**Fig. 1A** and **B**)^13–18^. Mammalian TRPA1 is a ligand-gated chemosensor that is activated by covalent modification of conserved cytoplasmic cysteine residues by environmental and endogenous electrophiles including allyl isothiocyanate (AITC), cinnamaldehyde, and acrolein (**Fig. 1A**, grey circles)^19–22^. In sensory neurons, active TRPA1 channels then facilitate extracellular Ca^2+^ entry to initiate pain signals and neuropeptide release^13,23–28^. Neuropeptide-evoked immune responses release pro-inflammatory mediators that sensitize neurons to subsequent painful stimuli by activating or priming receptors including TRPA1^13,29–31^. The ability of TRPA1 to initiate pain and neurogenic inflammation as well as be sensitized by these signals places it in the center of a regulatory pathway that could go awry^13^. Accordingly, animal knockout studies show that TRPA1 plays a key role in the establishment of neurogenic inflammation and neuronal hypersensitivity making it a prime drug target for managing chronic pain and inflammation^15,32–34^.

**Figure 1.**
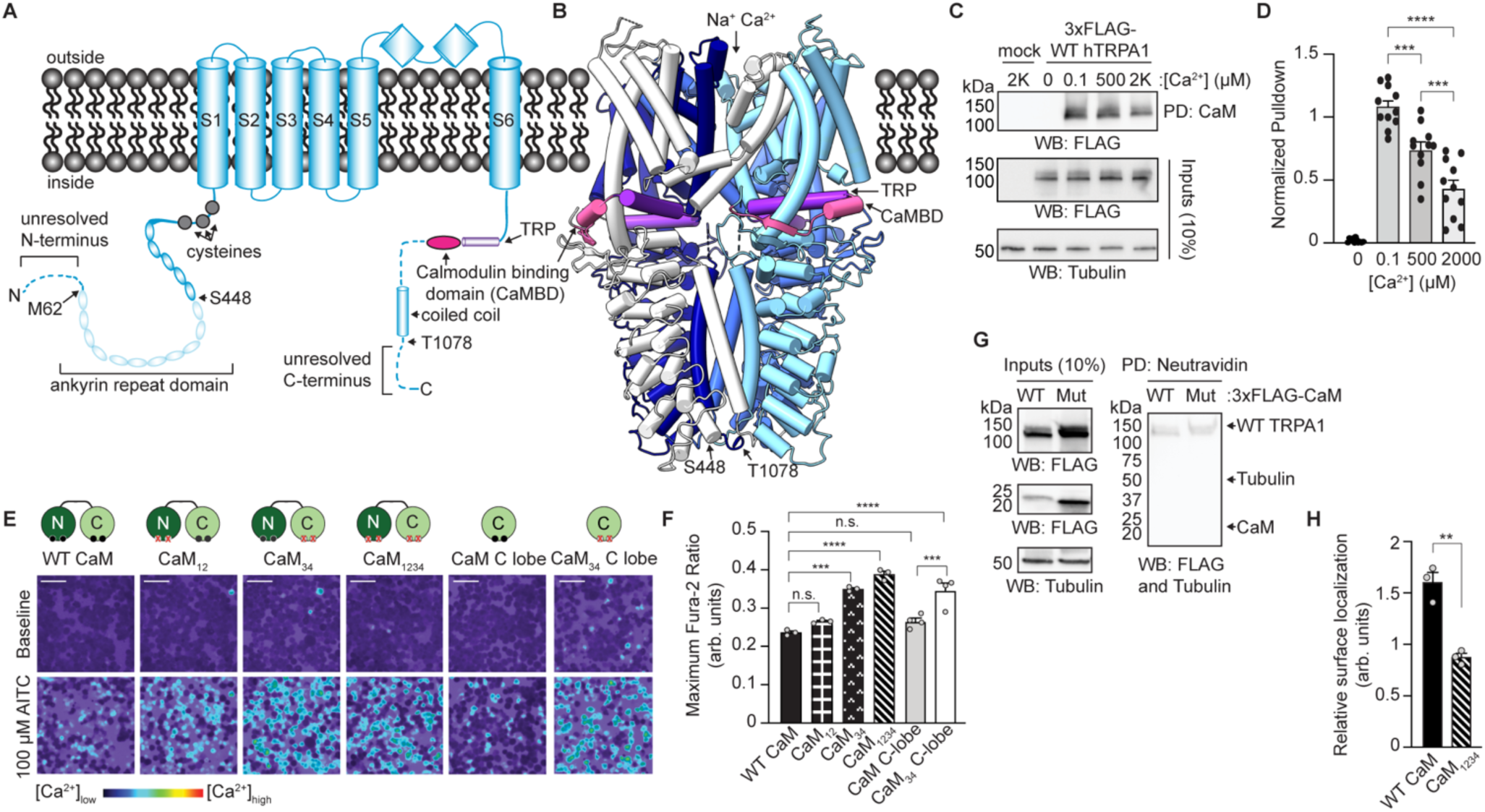
Calmodulin (CaM) C-lobe binding regulates TRPA1 activity. (**A**) Cartoon schematic of a full-length hTRPA1 monomeric subunit with relevant structural features and a previously identified Calmodulin binding domain (CaMBD, pink) denoted. Dashes and transparencies indicate unresolved regions in Cryo-EM structures. Residues denoted with arrows indicate positions of truncations used in this study. (**B**) Ribbon diagram of WT hTRPA1 atomic model for residues S448- T1078 from the homotetrameric channel (PDB: 6V9W). Each subunit is colored differently for clarity. The CaMBD and TRP helix are colored as in A. (**C**) WT TRPA1 interacts with CaM in a Ca^2+^-dependent manner. Immunoblotting analysis of 3xFLAG-WT hTRPA1 after CaM-agarose pulldown at the indicated Ca^2+^ concentrations from lysates of HEK293T cells transfected with 3xFLAG-WT hTRPA1 or empty vector (mock). Samples were probed using an HRP-conjugated anti-FLAG antibody. Tubulin from whole cell lysates (10%, inputs) was the loading control. (**D**) Quantitative analysis of CaM-agarose enrichment of 3xFLAG-WT hTRPA1 at the indicated Ca^2+^ concentrations relative to the maximum enrichment within each replicate. Data represent mean ± SEM. ***p=0.0003 (0.1 versus 500 µM Ca^2+^), ***p=0.0008 (500 versus 2000 µM Ca^2+^), ****p<0.0001 (0.1 versus 2000 µM Ca^2+^). n=11, one-way ANOVA with Tukey’s *post hoc* analysis. (**E**) Co-expression with a functional CaM C-lobe suppresses WT TRPA1 activity. Ratiometric Ca^2+^ imaging of HEK293T cells transfected with WT hTRPA1 and WT or the indicated CaM mutants. Cells were stimulated with 100 µM AITC. Images are representative of three (WT CaM, CaM_12_, CaM_34_, and CaM_1234_) or four (CaM C-lobe and CaM_34_ C-lobe) independent experiments. Scale bars indicate 50 µm. (**F**) Quantification of 100 µM AITC-evoked change in Fura-2 ratio of data from panel (E). Data represent mean ± SEM. ****, p<0.0001, ***p=0.001 (WT CaM versus CaM_34_) or 0.0007 (CaM C-lobe versus CaM_34_ C-lobe), n.s. not significant p>0.05. n= 3 (WT CaM, CaM_12_, CaM_34_, and CaM_1234_) or 4 (CaM C-lobe and CaM_34_ C-lobe) independent replicates, one-way ANOVA with Tukey’s *post hoc* analysis. (**G**) Surface biotinylation. Immunoblotting analysis of 3xFLAG-WT hTRPA1, 3xFLAG-WT CaM or 3xFLAG-CaM_1234_ protein expression in biotin-labeled plasma membranes from transfected cells. Biotinylated proteins were precipitated by Neutravidin resin pulldown and probed as in Fig. 1C. CaM and Tubulin were the negative controls for plasma membrane localization. (**H**) Quantitative analysis of the plasma membrane localization of 3xFLAG-WT hTRPA1 relative to Tubulin inputs. Data represent mean ± SEM. **p=0.0025, n=3, two-tailed Student’s t-test.

TRPA1 exhibits complex Ca^2+^ regulation; Ca^2+^ entry initially enhances channel activity (*e.g.*, potentiation), but then causes rapid inactivation (*e.g.*, desensitization) as cytoplasmic Ca^2+^ rises^21,35,36^. TRPA1 electrophile agonist modifications can persist for at least 10 minutes, yet TRPA1 desensitizes on a millisecond timescale^35,37^. Thus, Ca^2+^ regulation is critical to limit the TRPA1 functional window, however, the mechanism underlying this regulation is poorly understood. Previous work suggests that TRPA1 potentiation and desensitization are independent regulatory events that may engage distinct channel machinery^36,38^. Direct Ca^2+^ binding sites have been proposed in TRPA1 cytoplasmic and transmembrane domains to control potentiation and/or desensitization, however, deep mechanistic insight is lacking to explain whether and how these sites affect Ca^2+^ regulation and many of these sites remain controversial (**Fig. 1A** and **B**) ^38–40^. Moreover, most of these elements reside within or near key TRPA1 structural domains that contribute to channel gating. Genetic perturbation in these elements might compromise intrinsic channel structure or function, further complicating interpretation of their effects on Ca^2+^ regulation (**Fig. 1A** and **B**).

Ion channels can also be regulated by the universal Ca^2+^ sensor Calmodulin (CaM), which was recently shown to bind TRPA1 in a Ca^2+^-dependent manner and affect its Ca^2+^ regulation^21,36,39–41^. CaM was proposed to mediate these effects in part through a CaM binding domain (CaMBD) located adjacent to the TRP helix in the membrane proximal TRPA1 cytoplasmic C-terminus (**Fig. 1A** and **B**, CaMBD in pink and TRP helix in purple)^41^. Genetic perturbation of the CaMBD had modest effects on TRPA1 Ca^2+^ regulation perhaps due to incomplete loss of CaM binding, which raised the intriguing possibility that TRPA1 contains another CaM binding site^41^. Here, we identify a previously unreported, highly conserved, high-affinity <u>CaM</u> <u>b</u>inding <u>e</u>lement in the <u>d</u>istal structurally unresolved TRPA1 <u>C</u>-terminus (the DCTCaMBE) that we propose to be the main site for a TRPA1:Ca^2+^/CaM interaction. We show that TRPA1 binds CaM best at basal Ca^2+^ concentration through its DCTCaMBE and that this interaction can be ablated with short truncations, single point mutations, or a TRPA1 C-terminal peptide without affecting Ca^2+^-independent channel properties, allowing us to decouple Ca^2+^/CaM binding and TRPA1 function for the first time. TRPA1 DCTCaMBE mutants revealed that Ca^2+^/CaM binding to this site is dispensable for proper potentiation, but it is critically required for rapid desensitization providing further support for mechanistic independence of these two regulatory events. To date, most CaM regulated ion channels engage both the N- and C-lobes wherein Ca^2+^-mediated conformational changes in CaM confer Ca^2+^ sensing to the regulated channel^42–59^. In contrast, we find that the TRPA1 DCTCaMBE exclusively binds the Ca^2+^/CaM C-lobe and that only the CaM C-lobe was necessary for TRPA1 rapid desensitization. Thus, we propose that CaM binding *per se* likely does not directly trigger desensitization. Instead, we found desensitization resistant TRPA1 channels could be partially rescued by increasing the extracellular Ca^2+^ concentration available to permeate through open channels. Thus, we present a model that Ca^2+^/CaM acts as a long-range allosteric regulator to prime an intrinsic TRPA1 Ca^2+^ binding site that is the true desensitization gate. In this way, we propose that Ca^2+^/CaM serves as a regulatory binding partner for TRPA1 akin to the auxiliary subunits of voltage-gated ion channels to establish the proper TRPA1 functional window in cells^60–64^.

## RESULTS

### Ca^2+^/Calmodulin regulates TRPA1 in cells via its C-lobe

To initially ask whether CaM can directly regulate TRPA1, we assayed wild type (WT) human TRPA1 (hTRPA1) for its ability to bind CaM-agarose at distinct Ca^2+^ concentrations. Consistent with literature precedence^41^, hTRPA1 failed to bind CaM in the absence of Ca^2+^ (**Fig. 1C** and **D**, 0 µM Ca^2+^). Instead, we observed a robust interaction with hTRPA1 at resting, cytosolic Ca^2+^ concentration (0.1 µM) with a stepwise decrease in binding at intermediate (500 µM) and extracellular (2 mM) Ca^2+^ concentrations (**Fig. 1C** and **D**). These results are consistent with prior results that TRPA1 binds Ca^2+^/CaM, but not apo CaM^41^.

We next probed whether CaM regulates TRPA1 function by co-expressing WT hTRPA1 in HEK293T cells with WT CaM or a CaM variant that has a point mutation in each EF hand that prevents Ca^2+^ binding (CaM_1234_), which was previously shown to be unable to bind to TRPA1^41^, and monitoring channel activity by ratiometric Ca^2+^ imaging. Cells co-expressing WT CaM exhibited reduced AITC-evoked hTRPA1 activity at both sub-saturating and saturating agonist concentrations compared to cells co-expressing CaM_1234_ (**Fig. 1E** and **F**, solid black versus striped). These differences in channel activity were not due to WT or mutant CaM-induced changes to hTRPA1 expression or subcellular localization (**Fig. 1G, H** and **S1A**).

Previous work showed that co-expression with WT CaM or a CaM mutant with only a functional C-lobe (CaM_12_) enhanced mouse TRPA1 channel desensitization, demonstrating that the CaM C-lobe is critical to this regulation^41^. In our ratiometric Ca^2+^ imaging assay, enhanced desensitization would appear as reduced channel activity. To ask which CaM lobes are important to our observed activity suppression, we co-expressed WT hTRPA1 with different CaM constructs. Consistent with prior electrophysiological recordings^41^, we found that CaM_12_ significantly suppressed hTRPA1 activity, while a CaM mutant that only retained a functional N-lobe (CaM_34_) did not (**Fig. 1E** and **F**, black boxed versus carat bars). These results hint that the CaM C-lobe is sufficient to mediate hTRPA1 regulation; however, it is possible that the CaM N-lobe, independent of Ca^2+^ binding competency, is also required. Thus, we co-expressed the isolated CaM C-lobe or a Ca^2+^-binding deficient mutant (CaM_34_ C-lobe) with WT hTRPA1 and assayed for channel activity. Such experiments revealed that the CaM C-lobe suppressed hTRPA1 activity akin to WT CaM and CaM_12_, while the CaM_34_ C-lobe mutant did not (**Fig. 1E** and **F**, gray versus white bars). Importantly, none of the CaM constructs affected hTRPA1 expression (**Fig. S1A**).

Collectively, these results suggest that CaM suppresses WT hTRPA1 activity predominantly through its C-lobe. This recapitulates and expands on prior electrophysiological data showing that the CaM C-lobe is necessary for TRPA1 regulation^41^. While HEK293T cells endogenously express WT CaM, a large majority is likely pre-bound to other effector proteins leaving low levels of free CaM^65^. hTRPA1 overexpression seemingly overwhelms this free CaM pool yielding enhanced channel activity, which can be rescued by concurrent overexpression of WT CaM (**Fig. 1E**). Moving forward, we sought to identify the site and mode of Ca^2+^/CaM binding as well as the role for Ca^2+^/CaM binding in TRPA1 regulation.

### The TRPA1 distal C-terminus contains a critical Ca^2+^/Calmodulin binding element

A Ca^2+^/CaM binding domain (CaMBD) was previously identified near the TRP helix, which is involved in channel gating and resides within the structurally resolved core of hTRPA1 (**Fig. 1A** and **B**, pink)^41^. While the CaMBD was proposed to contribute to TRPA1 Ca^2+^ regulation, mutagenesis of this site did not ablate Ca^2+^/CaM binding and suggested that TRPA1 contains another Ca^2+^/CaM binding element^41^. To ask whether and to what extent the CaMBD governs the hTRPA1-Ca^2+^/CaM interaction, we tested a minimal hTRPA1 construct that was previously used for structural determination^66^ and which retains the CaMBD (hTRPA1^S488-T1078^) for CaM binding (**Fig. 1A, B** and **S2**). However, this minimal hTRPA1 construct failed to bind CaM *in vitro* indicating that the CaMBD is not sufficient for mediating this interaction (**Fig. 2A**).

**Figure 2.**
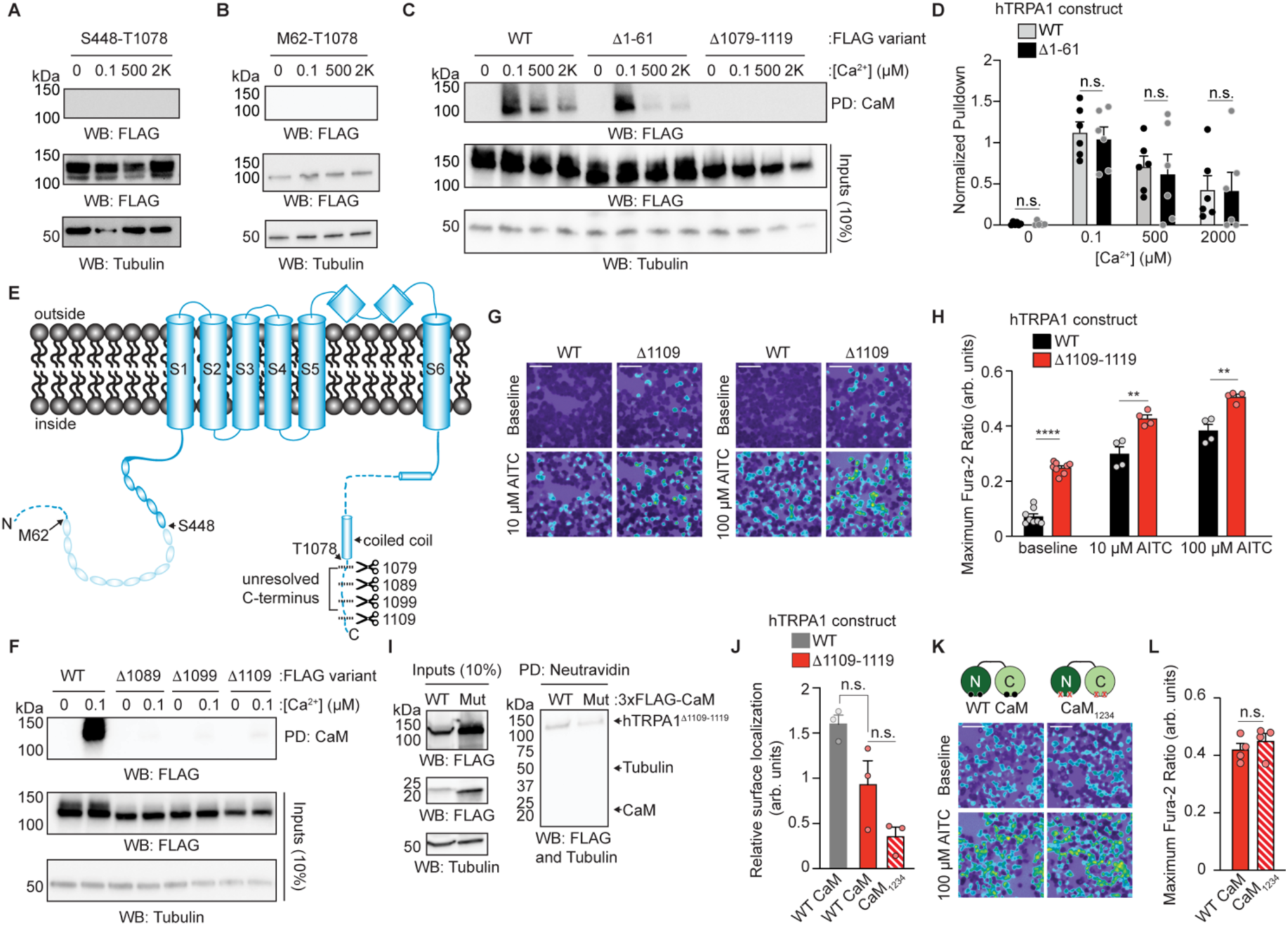
Identification of a Ca^2+^/CaM binding element in the TRPA1 distal C-terminus. (**A**-**C, F**) Representative immunoblotting analysis of 3xFLAG-tagged WT or truncation hTRPA1 constructs after CaM-agarose pulldown at the indicated Ca^2+^ concentrations from lysates of transfected HEK293T cells. Samples were probed as in Fig. 1C. Data are representative of three (A, B, and F) or six (C) independent experiments. Truncations in A-C are diagramed in Fig. 1A. (**D**) Quantitative analysis of CaM-agarose enrichment of 3xFLAG-WT hTRPA1 (grey) or 3xFLAG-hTRPA1^Δ1–61^ (black) at the indicated Ca^2+^ concentrations relative to the maximum enrichment within each replicate. Data represent mean ± SEM. n.s. not significant p>0.05, n=6, two-tailed Student’s t-test. (**E**) Simplified cartoon schematic from Fig. 1A denoting sites of truncations used in A-D and a suite of hTRPA1 unresolved C-terminus truncations tested in F-L. (**G**) Ratiometric Ca^2+^ imaging of HEK293T cells transfected with 3xFLAG-WT hTRPA1 or 3xFLAG-hTRPA1^Δ1109-1119^. Cells were stimulated with AITC (10 or 100 µM). Images are representative of four independent experiments. Scale bars indicate 50 µm. (**H**) Quantification of baseline, 10 µM or 100 µM AITC-evoked changes in Fura-2 ratio from cells in (G) transfected with WT hTRPA1 (black) or hTRPA1^Δ1109-1119^ (red). Data represent mean ± SEM. ****p<0.0001 (baseline), **p=0.0043 (10 µM AITC), and **p=0.0025 (100 µM AITC). n = 4 independent experiments, n ≥ 90 cells per transfection condition per experiment, two-tailed Student’s t-test. (**I**) Surface biotinylation. Immunoblotting analysis of 3xFLAG-hTRPA1^Δ1109-1119^, 3xFLAG-WT CaM or 3xFLAG-CaM_1234_ protein expression in biotin-labeled plasma membranes from transfected cells. Biotinylated proteins were precipitated by Neutravidin resin pulldown and probed as in Fig. 1G. CaM and Tubulin were the negative controls for plasma membrane localization. (**J**) Quantitative analysis of the plasma membrane localization of 3xFLAG-WT hTRPA1 (grey) or 3xFLAG-hTRPA1^Δ1109-1119^ (red) relative to Tubulin. WT hTRPA1 quantification is repeated from Fig. 1H. Data represent mean ± SEM. n.s. not significant p>0.05, n=3, one-way ANOVA with Bonferroni’s *post hoc* analysis. (**K**) Ratiometric Ca^2+^ imaging of HEK293T cells co-transfected with 3xFLAG-hTRPA1^Δ1109-1119^ and 3xFLAG-WT CaM or CaM_1234_. Cells were stimulated with 100 µM AITC. Images are representative of four independent experiments. Scale bars indicate 50 µm. (**L**) Quantification of 100 µM AITC-evoked change in Fura-2 ratio of data from panels (K) of cells co-expressing hTRPA1^Δ1109-1119^ and WT CaM (red bars) or CaM_1234_ (striped bars). Data represent mean ± SEM. n.s. not significant (p>0.05). n = 4 independent experiments, n ≥ 90 cells per transfection condition per experiment, two-tailed Student’s t-test.

The hTRPA1^S488-T1078^ construct truncates the disordered N-terminus (amino acids 1-61), the structurally unresolved ankyrin repeats (ARs, amino acids 61-447 containing AR1-11), and the structurally unresolved distal C-terminus (amino acids 1079-1119) suggesting one or more of these domains contain critical CaM binding sites (**Fig. 1A**, dashed and transparent regions). TRPA1 is unique among the mammalian TRP channels in that it has the largest N-terminal AR domain (ARD) consisting of 16 ARs (**Fig. 1A**)^17,67^. Though a functional or regulatory role for the large TRPA1 ARD is unclear, ARs are known to mediate protein-protein interactions raising the possibility that the TRPA1 ARD contains a CaM binding site^68^. Surprisingly, we found that a hTRPA1 construct containing the full ARD but lacking the structurally unresolved N- and C-termini (hTRPA1^M62-T1078^) failed to bind CaM indicating that a critical CaM binding element resides in one or both of those regions (**Fig. 2B**). Thus, we made hTRPA1 truncations lacking only the unresolved N- or C-termini (hTRPA1^Δ1-61^ and hTRPA1^Δ1079-1119^, respectively) and found that while loss of the unresolved N-terminus did not affect Ca^2+^/CaM binding compared to WT hTRPA1, loss of the unresolved C-terminus ablated Ca^2+^/CaM binding (**Fig. 2C** and **D**).

The hTRPA1 unresolved C-terminus includes the terminal 41 amino acids, which accounts for only 3.7% of the channel sequence. An acidic cluster in this domain proximal to the coiled coil was previously proposed to be involved in Ca^2+^ regulation indicating a role for the unresolved C-terminus in channel modulation^38^. To initially map the distal C-terminal CaM binding element (DCTCaMBE), we built a suite of truncations lacking sequential blocks of 10 residues (**Fig. 2E**) and assayed them for CaM binding. We found that even the smallest of these truncations lacking only the terminal 11 amino acids (hTRPA1^Δ1109-1119^) failed to bind CaM at basal Ca^2+^ concentration (**Fig. 2F**) indicating that at least part of the DCTCaMBE resides within the extreme end of the hTRPA1 C-terminus.

### Loss of Ca^2+^/Calmodulin binding is associated with hyperactive TRPA1 channels

Identification of the DCTCaMBE in the distal C-terminus far from the hTRPA1 core structure raises the intriguing possibility that we could selectively ablate Ca^2+^/CaM binding without compromising intrinsic channel function. Such a construct would facilitate a detailed characterization of whether and how CaM regulates hTRPA1 without relying on CaM mutants or inhibitors that may disrupt other aspects of CaM biology or that could indirectly impact channel function. Excitingly, the hTRPA1^Δ1109-1119^ construct yielded functional channels with increased basal and agonist-evoked activity compared to WT hTRPA1 (**Fig. 2G** and **H**). This increased activity was not due to enhanced mutant protein production or surface localization (**Fig. 2I** and **J**) and instead suggests that loss of Ca^2+^/CaM binding yields hyperactive hTRPA1 channels.

To directly test whether the hTRPA1^Δ1109-1119^ construct relieves Ca^2+^/CaM-mediated channel suppression, hTRPA1^Δ1109-1119^ was co-expressed in HEK293T cells with WT CaM or CaM_1234_. Unlike WT hTRPA1 (**Fig. 1E** and **F**), we found that hTRPA1^Δ1109-1119^ channels were unaffected by CaM co-expression (**Fig. 2K** and **L**) consistent with a loss of Ca^2+^/CaM binding capability (**Fig. 2F**). Akin to WT channels, WT CaM and CaM_1234_ had no effect on hTRPA1^Δ1109-1119^ expression or surface localization (**Fig. 2I, J**, and **S1**). Together, these results suggest that the hTRPA1 distal C-terminus contains a critical DCTCaMBE that is required for Ca^2+^/CaM binding and its negative regulation of channel activity.

### TRPA1 and Calmodulin co-localize in cells

The pronounced Ca^2+^/CaM binding we observed at basal Ca^2+^ concentration (**Fig. 1C** and **D**, 100 nM Ca^2+^) raises the intriguing possibility that WT hTRPA1 and Ca^2+^/CaM form a stable complex at rest and that this interaction is lost in the hTRPA1^Δ1109-1119^ mutant. Consistently, deconvolved immunofluorescence (IF) images of Neuro2A cells co-expressing V5-tagged WT CaM and 3xFLAG-tagged WT hTRPA1 exhibited pronounced co-localization of WT hTRPA1 and WT CaM in resting cells (**Fig. 3A**, left). In contrast, the 3xFLAG-tagged hTRPA1^Δ1109-1119^ channels showed greatly reduced co-localization with WT CaM in Neuro2A cells (**Fig. 3A**, right). To further assess the degree of co-localization between CaM and WT hTRPA1 or the hTRPA1^Δ1109-1119^ mutant, we quantified the V5-CaM and 3xFLAG-TRPA1 intensities at every pixel in individual cells and calculated Pearson’s correlation coefficients (R) (**Fig. S3**)^69^. These results revealed a significant reduction in the CaM:TRPA1 co-localization with the hTRPA1^Δ1109-1119^ mutant compared to WT channels (**Fig. 3B**), suggesting that the loss of Ca^2+^/CaM binding in pulldown experiments translates to a significant disruption of the relative spatial distribution of TRPA1 and CaM in a cellular environment due to reduced protein-protein interaction.

**Figure 3.**
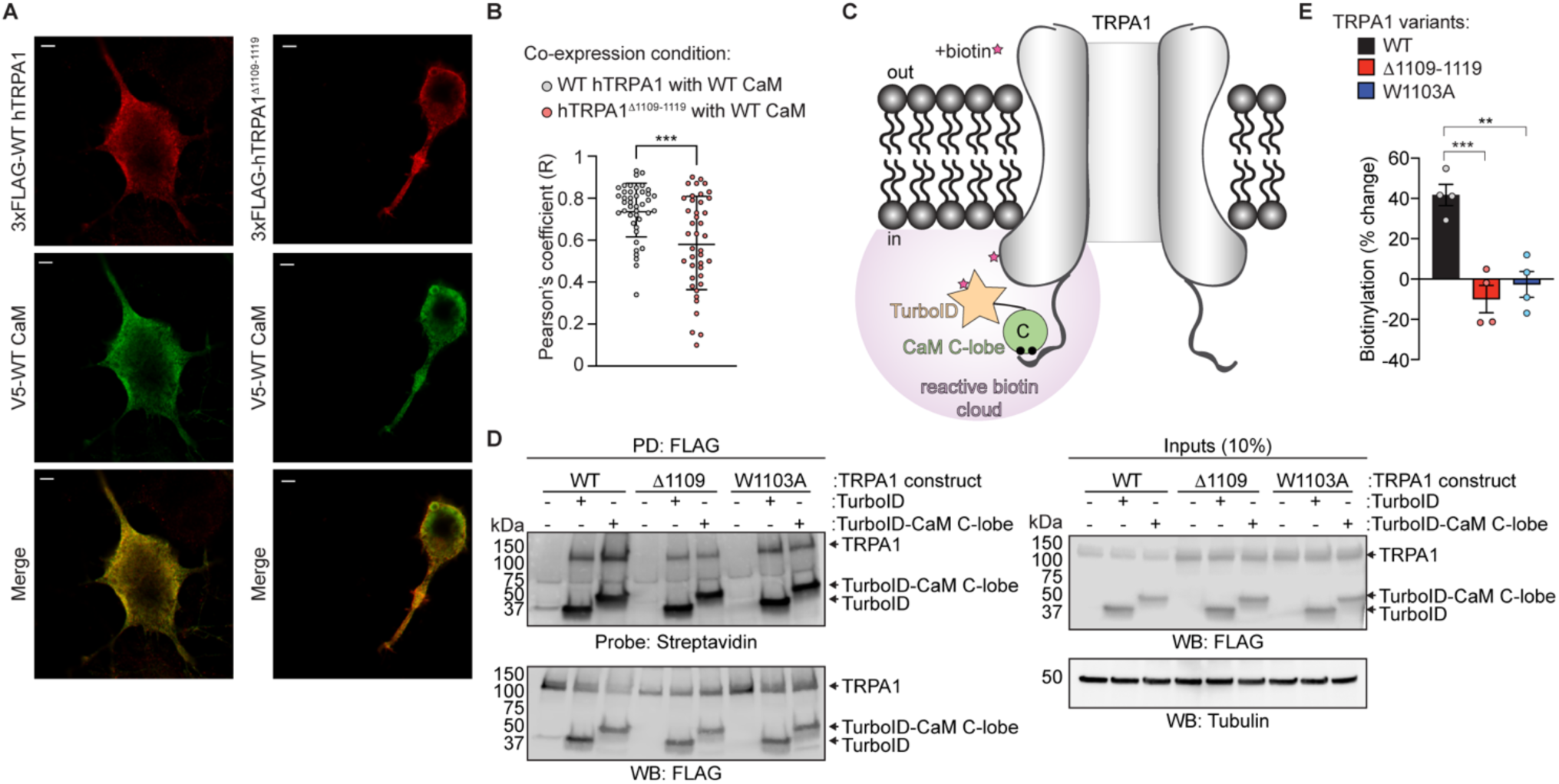
TRPA1 and CaM co-localize in cells. (**A**) Representative deconvolved Airyscan images of Neuro2A cells co-expressing 3xFLAG-WT hTRPA1 (left) or hTRPA1^Δ1109-1119^ (right) with V5-WT CaM. Cells were stained with anti-FLAG (red) and anti-V5 (green) antibodies. Scale bar indicates 2 µm. Images are representative of 3 independent experiments. (**B**) Pearson’s correlation coefficients (R) were determined for WT hTRPA1:WT CaM or hTRPA1^Δ1109-1119^:WT CaM using raw images (see Fig. S3). Data represent mean ± SD. ***p=0.0002, n=41 (WT) or 40 (Δ1109-1119) cells from 3 independent experiments, two-tailed Student’s t-test. n >10,000 pixels in 1 cell per condition. (**C**) Proximity biotinylation approach. TurboID (orange star) fused to the CaM C-lobe (green circle) is co-expressed with TRPA1 variants (black) in cells. Addition of biotin (pink star) to the media will facilitate generation of a local reactive biotin cloud (purple) that will biotinylate TurboID- CaM C-lobe and TRPA1, pending an interaction. (**D**) Immunoblotting analysis of biotinylated 3xFLAG-WT hTRPA1, hTRPA1^Δ1109-1119^, or W1103A hTRPA1 co-expressed with empty vector (-), 3xFLAG-TurboID-CaM C-lobe, or 3xFLAG- TurboID in HEK293T cells. FLAG immunoprecipitated eluates were probed for biotinylation with Streptavidin-HRP. FLAG- tagged proteins were probed using HRP-conjugated anti-FLAG antibody. Tubulin was the loading control. Blots representative of four independent experiments. (**E**) Quantification of the percent change to TRPA1 biotinylation with TurboID-CaM C-lobe versus TurboID from experiments as in (D). Pulldown Streptavidin-HRP (*e.g.*, biotinylation) was normalized to FLAG and Tubulin from inputs for each sample. Biotinylation by TurboID was subtracted from biotinylation by TurboID-CaM C-lobe. Data are presented as the percent change from biotinylation with TurboID. Data represent mean ± SEM. ***p=0.0005, **p=0.0013. n = 4 independent experiments, one-way ANOVA with Bonferroni’s *post hoc* analysis.

We next employed a proximity biotinylation assay to determine whether a direct protein-protein interaction between WT hTRPA1 and CaM occurs in intact cells at basal Ca^2+^ levels and whether this interaction is lost with the hTRPA1^Δ1109-1119^ mutant^70^. TurboID-fused CaM C-lobe or free TurboID tag were co-expressed with WT hTRPA1 or the hTRPA1^Δ1109-1119^ mutant and biotinylated proteins were detected in FLAG immunoprecipitation eluates (**Fig. 3C**). These experiments revealed that both TRPA1 constructs were biotinylated by the free TurboID tag, consistent with TRPA1 and cytoplasmic TurboID proteins occupying similar subcellular spaces (**Fig. 3D**). In contrast, WT hTRPA1 exhibited an approximately 40% increase in biotinylation when co-expressed with the TurboID-fused CaM C-lobe construct suggesting a direct interaction in cells (**Fig. 3D** and **E**). Interestingly, the hTRPA1^Δ1109-1119^ mutant exhibited reduced biotinylation with the TurboID-fused CaM C-lobe, which may be due to sequestration of TurboID-tagged CaM C-lobe by endogenous CaM effector proteins that would limit the stochastic interaction of TRPA1 with cytoplasmic TurboID (**Fig. 3D** and **E**). Collectively, these results support the model that WT TRPA1 and Ca^2+^/CaM associate at basal Ca^2+^ concentrations in cells.

### The TRPA1 distal C-terminus has a high affinity for the Ca^2+^/Calmodulin C-lobe

We next sought to quantify the affinity of the hTRPA1-Ca^2+^/CaM interaction. While quantitative binding assays with membrane proteins are particularly challenging, we developed hTRPA1 distal C-terminus peptides for such analyses. Maltose binding protein (MBP)-tagged versions of the full hTRPA1 unresolved C-terminus (hTRPA1^1079-1119^) and smaller fragments (hTRPA1^1089-1119^ and hTRPA1^1099-1119^) were expressed in HEK293T cells and tested for their ability to bind Ca^2+^/CaM. Each bound CaM in a Ca^2+^-dependent manner with no binding observed with free MBP tag, confirming that the C-terminus contains a DCTCaMBE (**Fig. S4A**). The hTRPA1^1089-1119^ peptide was selected for binding studies as it afforded the highest-yield bacterial purifications (**Fig. S4B**). Importantly, the purified hTRPA1^1089-1119^ peptide competed with WT hTRPA1 for Ca^2+^/CaM binding confirming that the isolated element can adopt a conformation amenable to Ca^2+^/CaM binding after bacterial purification and liberation from MBP (**Fig. S4C** and **D**).

To initially characterize the hTRPA1 DCTCaMBE:Ca^2+^/CaM interaction, we performed size exclusion chromatography (SEC)-based binding assays with the hTRPA1^1089-1119^ peptide and a suite of purified CaM constructs (**Fig. S4E-H**). These assays were performed by mixing equimolar amounts (100 µM) of peptide and CaM construct in the presence of 2 mM free Ca^2+^ to promote uniform complex formation and to simplify data interpretation. While hTRPA1^1089-1119^ and WT CaM eluted as spatially separate peaks when analyzed independently (**Fig. S5A**, blue and yellow traces), mixed hTRPA1^1089-1119^ and WT CaM eluted as a single peak slightly larger than free WT CaM with higher 280 nm absorbance indicating all available hTRPA1^1089-1119^ peptide was bound in a stable complex with CaM (**Fig. 4A** and **S4A,** green trace). This complex was disrupted by chelating Ca^2+^ with 5 mM EGTA, confirming the Ca^2+^-dependence of the interaction (**Fig. 4A**, black trace). Moreover, the Ca^2+^-insensitive CaM_1234_ mutant failed to bind the TRPA1 peptide consistent with no interaction between apo CaM and the TRPA1 distal C-terminus (**Fig. S5B**, red trace). Fractions from the SEC runs revealed co-elution of the hTRPA1^1089-1119^ peptide with WT CaM in the presence, but not the absence of Ca^2+^ (**Fig. 4A**, inset 2). To simulate a low Ca^2+^ environment^71^, we performed similar experiments with CaM_12_, which revealed that it also forms a complex with all available hTRPA1^1089-1119^ peptide only in the presence of Ca^2+^ (**Fig. 4B** and **S5C**, green versus black trace). These results were similarly supported by fractions from the SEC runs (**Fig. 4B**, inset 2). Our Ca^2+^ imaging experiments suggested that the CaM N-lobe is not required for TRPA1 regulation by Ca^2+^/CaM. To ask whether the N-lobe is required for CaM binding to the hTRPA1^1089-1119^ peptide, we performed SEC assays with the CaM C-lobe, which revealed that the CaM C-lobe can engage the hTRPA1^1089-1119^ peptide in isolation (**Fig. 4C**, green trace). These results suggest that neither the N-lobe nor Ca^2+^ binding in the N-lobe is required for the hTRPA1 DCTCaMBE-Ca^2+^/CaM interaction and that the CaM C-lobe is likely responsible for mediating the Ca^2+^-dependent binding to the hTRPA1 distal C-terminus.

**Figure 4.**
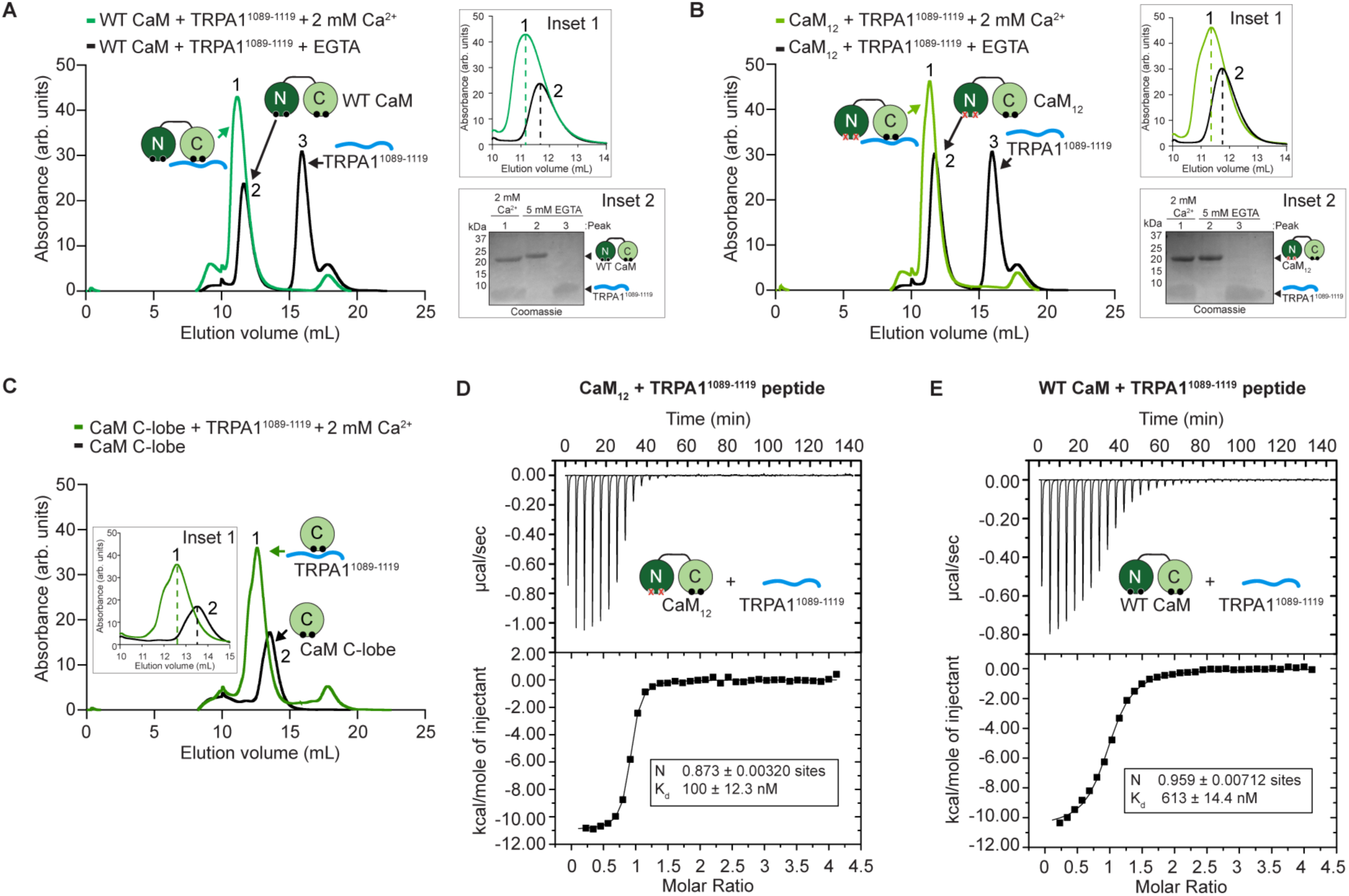
The TRPA1 distal C-terminus contains a high affinity Ca^2+^/CaM binding site. (**A** and **B**) Superdex 75 chromatograms of WT CaM (A) or CaM_12_ (B) with hTRPA1^1089-1119^ in the presence of 2 mM Ca^2+^ (green) or 5 mM EGTA (black). Inset 1 zooms in on the CaM and CaM:peptide peaks (peaks labeled 1 and 2). Inset 2 are Coomassie stains of fractions labeled 1 (green trace), 2 (black trace), and 3 (black trace). Arrows indicate CaM constructs and hTRPA1^1089-1119^ peptide. (**C**) Superdex 75 chromatograms of CaM C-lobe alone (black) or with hTRPA1^1089-1119^ in the presence of 2 mM Ca^2+^ (green). Chromatograms in A, B, and E were generated from 100 µM protein for each construct. Data are representative of three independent replicates. (**D** and **E**) The TRPA1 distal C-terminus has a higher affinity for CaM_12_ than WT CaM. Representative isothermal titration calorimetry plots of CaM_12_ (D) or WT CaM (E) titrated by hTRPA1^1089-1119^ peptide at 2 mM Ca^2+^ and fitted using the one-site binding model. Values for the number of binding sites (N) and the binding constant K_d_ are shown.

To quantitatively measure the hTRPA1^1089-1119^ peptide:Ca^2+^/CaM interaction we used isothermal titration calorimetry (ITC). As with the SEC assays above, these assays were performed with 2 mM Ca^2+^ to reduce heterogeneity and to simplify data interpretation. Low and high Ca^2+^ levels were simulated with CaM_12_ and WT CaM, respectively as reported previously^72,73^. Three independent runs of the TRPA1^1089-1119^ peptide with CaM_12_ and WT CaM revealed affinities centering on 100 nM and 613 nM, respectively (**Fig. 4D** and **E, S5D** and **E**). The ITC results were not due to titration artifacts from the peptide (**Fig. S5F**) and these binding affinities are within range of those reported for CaM-regulated ion channels^72,74–77^. The higher affinity of CaM_12_ than WT CaM for the hTRPA1^1089-1119^ peptide suggests that CaM undergoes a conformational change upon full calcification that reduces the binding affinity, which has been observed with some CaM complexes^78–80^. These affinities also provide insight into why we saw a reduced TRPA1-Ca^2+^/CaM interaction at higher Ca^2+^ concentrations in our CaM-agarose assays (**Fig. 1C** and **1D**). Notably, we only observed a single binding event with both CaM_12_ and WT CaM indicating that the CaM N-lobe does not also bind the effector to mediate a bridged interaction between adjacent subunits as shown previously for TRPV6^72^. Such lobe specificity has been observed for Ca^2+^/CaM binding sites in other ion channels^50,76,77,81^. Together with our IF imaging and proximity biotinylation data (**Fig. 3**), these results indicate that hTRPA1 is a physiologically relevant binding partner for CaM, that the Ca^2+^/CaM interaction is predominantly mediated by the C-lobe, and that hTRPA1 and Ca^2+^/CaM may pre-associate in cells prior to channel activation.

### NMR spectroscopy of the TRPA1 C-terminus with Calmodulin reveals a Ca^2+^/C-lobe binding mode

To initially determine how the hTRPA1 distal C-terminus binds CaM, we generated a model of the hTRPA1^1089-1119^ peptide in complex with WT CaM using AlphaFold2 Multimer^82,83^. This model predicts that CaM adopts an open, extended conformation as seen with other CaM complexes (**Fig. 5A** and **S6A**)^76,77,84,85^. Moreover, only the CaM C-lobe engages the peptide as an alpha helix in this model (**Fig. 5A**), consistent with the binding results above (**Fig. 1C**, **1D**, and **4**).

**Figure 5.**
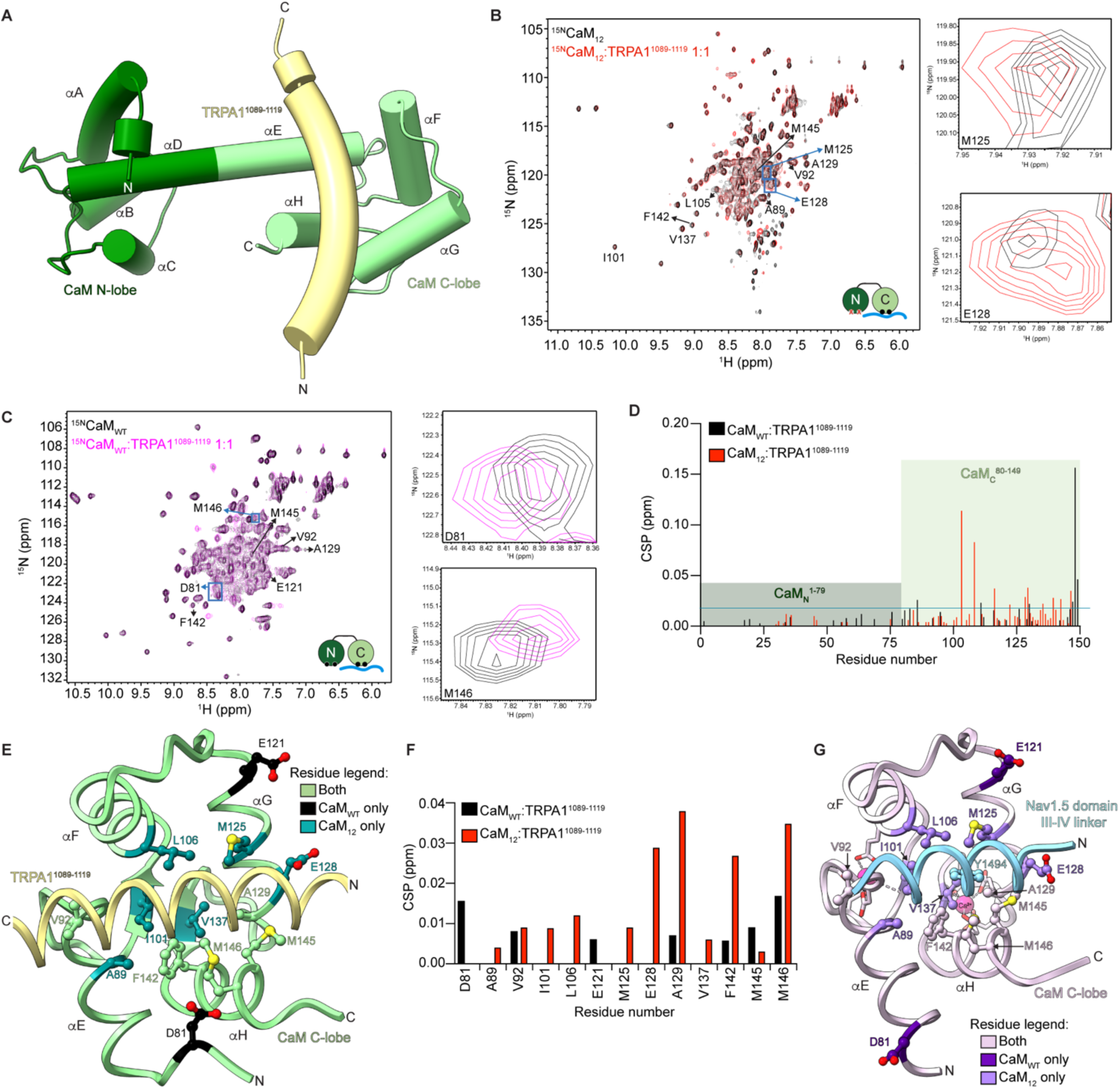
Analysis of the TRPA1 C-terminal tail Ca^2+^/CaM binding mode. (**A**) Ribbon diagram of the WT CaM atomic model (N-lobe, dark green, residues 1-79; C-lobe, light green, residues 80-149) in complex with part of the hTRPA1 C-terminus (yellow, residues 1089-1119) as predicted by AlphaFold2 Multimer. (**B**) Overlay of the ^15^N-^1^H HSQC spectra of ^15^N-labled CaM_12_ (black) with the ^15^N-labeled CaM_12_:TRPA1^1089-1119^ complex (red) at a 1:1 molar ratio. Insets depicts expanded view of the CSPs boxed in blue. (**C**) Overlay of the ^15^N-^1^H HSQC spectra of ^15^N-labled CaM_WT_ (black) with the ^15^N-labeled CaM_WT_:TRPA1^1089-1119^ complex (purple) at a 1:1 molar ratio. Insets depict expanded view of the CSPs boxed in blue. (**D**) CaM_12_ (red) and CaM_WT_ (black) chemical shift perturbations (CSPs) (δ_bound_ - δ_free_) as a function of residue number for the ^15^N-labeled CaM_12_:TRPA1^1089-1119^ or ^15^N-labeled CaM_WT_:TRPA1^1089-1119^ complex as in panels B and C, respectively. Dark and light green shadings denote CaM N- and C-lobe residues, respectively. The mean value plus one standard deviation is the horizontal blue line. (**E**) Ribbon diagram of the CaM C-lobe atomic model (green, residues 80-149) in complex with part of the hTRPA1 C-terminus (yellow, residues 1089-1119) as predicted by AlphaFold2 Multimer. Residues exhibiting CSPs from the ^15^N-labeled CaM_WT_:TRPA1^1089-1119^ complex (black), the ^15^N-labeled CaM_12_:TRPA1^1089-1119^ complex (teal), and both complexes (green) that face the TRPA1 C-terminus are depicted as balls and sticks. (**F**) CaM_WT_ (black) or CaM_12_ (red) CSPs for the C-lobe residues modeled in E from the ^15^N-labeled CaM_12_:TRPA1^1089-1119^ or CaM_WT_:TRPA1^1089-1119^ complexes from panels B and C, respectively. (**G**) Ribbon diagram of the crystal structure of the Ca^2+^/CaM:Nav1.5 DIII-IV linker complex (PDB entry 4DJC) (Sarhan et al PNAS **2012** 109: 3558-3563). The CaM C-lobe (purple), Nav1.5 peptide (blue), and Ca^2+^ ions (pink) are indicated. The CaM C-lobe residues exhibiting CSPs from the ^15^N-labeled CaM_WT_:TRPA1^1089-1119^ complex (dark purple), the ^15^N-labeled CaM_12_:TRPA1^1089-1119^ complex (medium purple), and both complexes (light purple) from panel E and F are depicted as balls and sticks. The key Nav1.5 interacting residue Y1494 is highlighted (blue).

We next empirically investigated the CaM lobe specificity of the Ca^2+^/CaM:TRPA1 C-terminal peptide interaction by collecting ^15^N-^1^H HSQC NMR spectra of ^15^N-labeled CaM_12_ or WT CaM alone and with unlabeled hTRPA1^1089-1119^ peptide at 1:1 ratios (**Fig. 5B** and **C**). Chemical shift perturbations (CSPs), which yield residue specific information regarding the binding interaction, were observed predominantly in the C-lobe for both CaM_12_ and WT CaM in the presence of the hTRPA1^1089-1119^ peptide, consistent with a C-lobe driven interaction (**Fig. 5D**, red and black, respectively). Moreover, CaM_12_ exhibited more CSPs in the C-lobe than WT CaM, consistent with a more engaged binding event (**Fig. 5D**, red versus black). To determine whether there is a conformational change between the CaM_12_:TRPA1 peptide and the WT CaM:TRPA1 peptide complexes that might explain the differences in binding affinities observed above, we subtracted the CSPs for CaM_12_:TRPA1^1089-1119^ (**Fig. 5B**) from those for CaM_WT_:TRPA1^1089-1119^ (**Fig. 5C**), which yields differential binding effects between the two complexes (**Fig. S6B**). This analysis was used previously to reveal CaM N-lobe engagement of a TRPV6 peptide with WT CaM^72^. Here, it revealed a loss of peaks for many C-lobe residues as well as some increased peaks at residues within the linker (residues 73-83), supporting a differential engagement of the TRPA1 peptide by the C-lobe, and only the C-lobe, in CaM_12_ versus WT CaM. This is likely due to a conformational change that may account for the weaker binding affinity in the fully calcified state.

To identify residues within our NMR spectra that may directly face the TRPA1 C-terminal peptide, we mapped the residues with CSPs from the CaM_12_:TRPA1^1089-1119^ and CaM_WT_:TRPA1^1089-1119^ spectra to an AlphaFold model of the hTRPA1^1089-1119^ peptide in complex with the CaM C-lobe (**Fig. 5E**). The CaM C-lobe model was used since it yielded a higher confidence model than that with WT CaM (**Fig. S6A** and **C**). This analysis identified five hydrophobic residues lining the CaM C-lobe hydrophobic pocket that showed chemical shifts in both the CaM_12_ and WT CaM spectra (**Fig. 5E**, green), an additional five hydrophobic residues and one acidic residue uniquely observed in the CaM_12_ spectra (**Fig. 5E**, teal), and two acidic residues uniquely observed in the WT CaM spectra (**Fig. 5E**, black). Analysis of CSPs for these peptide-facing residues reveals more and larger shifts from the CaM_12_:TRPA1^1089-1119^ complex than the CaM_WT_:TRPA1^1089-1119^ complex further supporting a more engaged binding event (**Fig. 5F**).

We used the Dali server to identify structures in the PDB with similarity to our hTRPA1^1089-1119^ peptide in complex with the CaM C-lobe model^86^. The top twenty results are for Ca^2+^/CaM:effector complexes where the calcified C-lobe engages the effector alone or in coordination with the calcified N-lobe (**Fig. S6D**). Alignment of our AlphaFold2 model with the Ca^2+^/CaM:Nav1.5 domain III-IV linker peptide crystal structure (PDB: 4DJC), in which only the C-lobe engages the peptide, showed a similar conformation for the CaM C-lobe and a similar binding mode to the effector peptides (**Fig. S6E**). Mapping the residues identified in our NMR spectroscopy experiments (**Fig. 5F**) to the Ca^2+^/CaM:Nav1.5 domain III-IV linker peptide crystal structure shows that the hydrophobic residues face the bound effector peptide (**Fig. 5G**). Collectively, these results suggest that Ca^2+^/CaM binds the TRPA1 distal C-terminus through its calcified C-lobe only and that further calcification of the N-lobe induces a conformational change that weakens the binding event.

### Mapping the highly conserved DCTCaMBE within the TRPA1 distal C-terminus

We next used our hTRPA1^1089-1119^ peptide in complex with the CaM C-lobe model with the residues implicated by our NMR spectroscopy studies to identify the hTRPA1 C-terminal residues that comprise the DCTCaMBE (**Fig. 6A**). Our model predicts that the CaM C-lobe binds to the hTRPA1 C-terminal alpha helix at a motif spanning amino acids 1099-1111 (**Fig. 6A**, goldenrod). The bound helix is predicted to adopt the same binding interface and bind in the same N-to-C direction as the Nav1.5 domain III-IV linker peptide (**Fig. S6E**). These residues forming the putative DCTCaMBE contain a hydrophobic ridge comprised of residues W1103, V1106, L1107, and V1110 which bind the hydrophobic cleft of the CaM C-lobe (**Fig. 6B**). This hydrophobic ridge is flanked on the N-terminal side by positively charged residues R1099 and R1102, and on the C-terminal side by K1111 with all three residues predicted to form salt bridges with glutamate residues from the CaM C-lobe (**Fig. 6C**). This putative DCTCaMBE is consistent with loss of CaM binding in the hTRPA1^Δ1109-1119^ truncation, which lacks the V1110 and K1111 residues (**Fig. 2E**).

**Figure 6.**
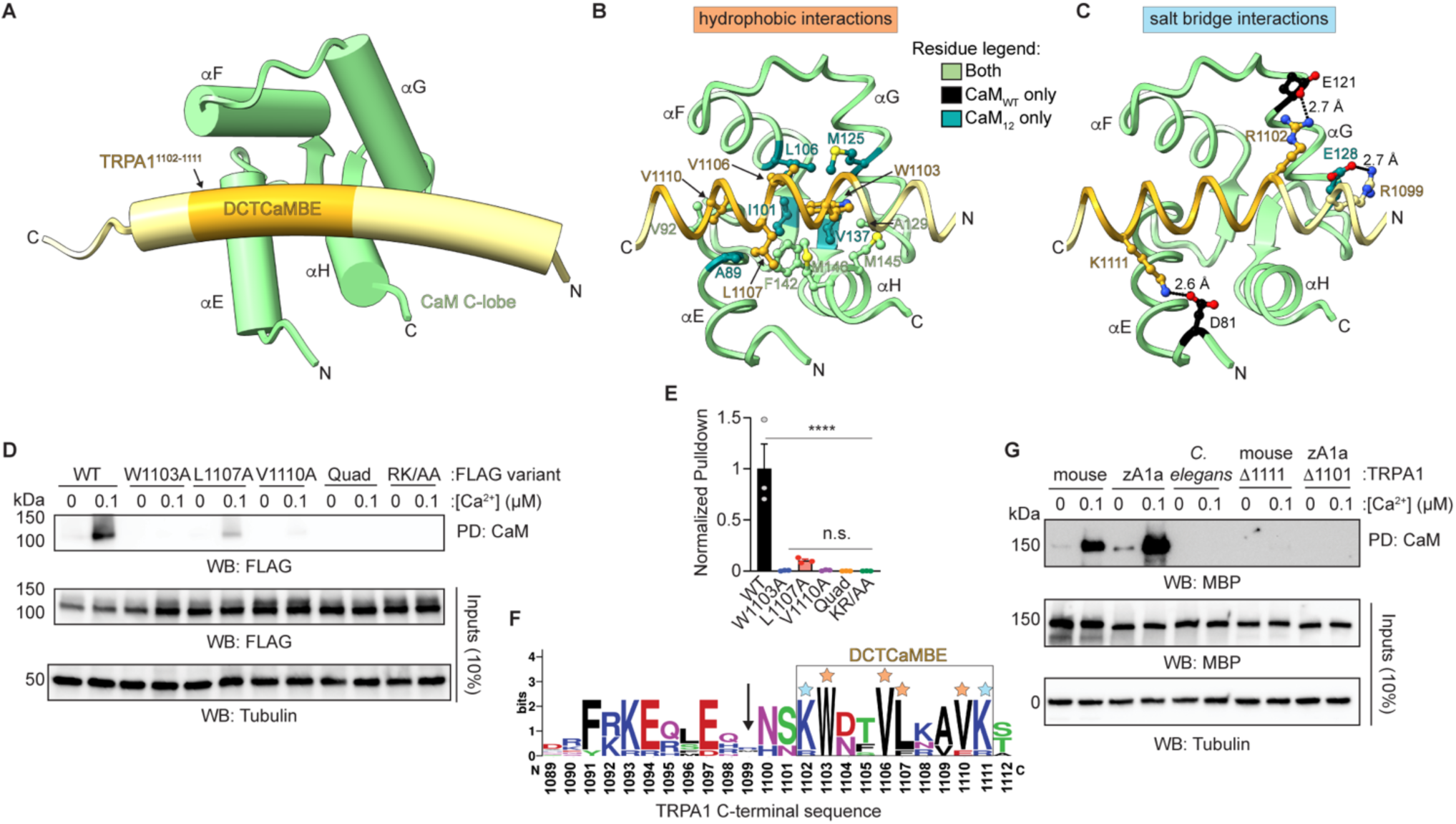
Identification of key and conserved TRPA1 DCTCaMBE residues involved in Ca^2+^/CaM binding. (**A**) (Right) Ribbon diagram of the CaM C-lobe atomic model (green, residues 80-149) in complex with part of the hTRPA1 C-terminus (yellow, residues 1089-1119) as predicted by AlphaFold2 Multimer. The region proposed to form the distal C-terminal CaM binding element (DCTCaMBE, hTRPA1 residues R1102-K1111) is indicated in goldenrod. (**B** and **C**) Ribbon diagrams with residues mediating hydrophobic interactions (B) or salt bridge interactions (C) depicted as balls and sticks. Residues colored as in Fig. 5E. The highlighted CaM residues exhibited CSPs in 2D NMR data sets (see Fig. 5F). (**D**) Immunoblotting analysis of the indicated 3xFLAG-hTRPA1 constructs after CaM-agarose pulldown in the absence or presence of Ca^2+^ from lysates of HEK293T cells transfected with 3xFLAG-WT, W1103A, L1107A, V1110A, W1103A/V1106A/L1107A/V1110A (Quad), or R1102A/K1111A (RK/AA) hTRPA1. Blot is representative of three independent experiments. Samples were probed as in Fig. 1C. (**E**) Quantification of CaM-agarose pulldowns represented in D. Pulldown was normalized to the WT hTRPA1 with Ca^2+^ average. Data represent mean ± SEM. ****p<0.0001, n.s. not significant (p>0.05). n = 3 independent experiments, one-way ANOVA with Tukey’s *post hoc* analysis. (**F**) Sequence alignment of nine TRPA1 orthologues aligned to residues 1089-1112 of hTRPA1. Alignment generated with Sequence Logo. Arrow denotes poorly conserved hTRPA1 R1099. Box denotes TRPA1 DCTCaMBE. Hydrophobic (orange) and salt bridge (blue) residues proposed to form the DCTCaMBE are denoted with stars. (**G**) Immunoblotting analysis of the indicated MBP-TRPA1 species orthologue constructs after CaM- agarose pulldown in the absence or presence of Ca^2+^ from lysates of HEK293T cells transfected with MBP-WT mouse, zebrafish TRPA1a isoform (zA1a), or *C. elegans* TRPA1 or the Δ1109-1119 equivalents of mouse (Δ1111-1125) or zebrafish (Δ1101-1115) TRPA1. Blot is representative of four independent experiments. Samples were probed using an anti-MBP primary antibody and an HRP-conjugated anti-mouse secondary antibody. Tubulin from whole cell lysates (10%, inputs) was the loading control.

To discern which of these residues comprise the DCTCaMBE, we built a suite of hTRPA1 alanine mutations and analyzed them for CaM binding. Individual mutation of W1103 or V1110 to alanine completely ablated CaM binding at basal Ca^2+^ concentration with a partial reduction in binding by a L1107A mutant (**Fig. 6D** and **E**). Mutating all four hydrophobic residues (Quad) or the flanking R1102 and K1111 residues (RK/AA) to alanine similarly ablated CaM binding at basal Ca^2+^ concentration (**Fig. 6D** and **E**). R1099 was not tested since it is poorly conserved (**Fig. 6F**, black arrow). These data support that R1102, W1103, V1106, L1107, V1110, and K1111 form the hTRPA1 DCTCaMBE, consistent with the structural prediction (**Fig. 6F**, box).

W1103 in the DCTCaMBE is predicted to adopt the same binding site as Y1494 which forms the main hydrophobic anchor in the Nav1.5 domain III-IV linker (**Fig. S6F**). Our model predicts that W1103 binds a hydrophobic pocket lined by residues identified in both the CaM_12_ and the WT CaM NMR data sets (A129, F142, M145, and M146) as well as residues uniquely identified in the CaM_12_ data set (I101, L106, M125, and V137) (**Fig. 6B**). L1107 is similarly located near residues identified in both NMR data sets (F142 and M145) and residues uniquely identified in the CaM_12_ data set (A89) while V1106 is near CaM_12_ unique residues (L106 and M125) and V1110 is near a residue identified in both (V92) (**Fig. 6B**). Many of the unique CaM_12_ residues that are near W1103 (I101, L106, and V137) are deep in the hydrophobic pocket and may hint at a wider C-lobe conformation binding the hTRPA1 DCTCaMBE at basal Ca^2+^ concentration. The WT CaM NMR data set uniquely identified D81 and E121, which are predicted to form salt bridges with K1111 and R1102 in the hTRPA1 DCTCaMBE, respectively while the CaM_12_ data set identified E128 that is predicted to interact with R1099 (**Fig. 5F** and **6C**). Though D81 and E121 were not identified in the CaM_12_ NMR data set, mutation of K1111 and R1102 in the hTRPA1 DCTCaMBE ablates Ca^2+^/CaM binding at basal Ca^2+^ concentration indicating that interactions with these residues are also important when cells are at rest (**Fig. 6D** and **E**). Future structural work is needed to capture the TRPA1 DCTCaMBE and Ca^2+^/CaM complex at distinct Ca^2+^ concentrations, however, our NMR data suggest that this interaction is predominantly driven by hydrophobic interactions at basal Ca^2+^ concentrations and by a combination of fewer hydrophobic interactions and salt bridge interactions at higher Ca^2+^ concentrations.

Prior work demonstrated that CaM regulates the mouse TRPA1 channel^41^. The hTRPA1 DCTCaMBE identified in this study is highly conserved among vertebrate TRPA1 including the mouse orthologue, but it is absent from *C. elegans* TRPA1 (**Fig. 6F** and **S6G**). To ask whether the DCTCaMBE governs CaM binding in other TRPA1 species orthologues, we assayed the CaM binding ability of WT and partial DCTCaMBE truncations of mouse, zebrafish, and *C. elegans* TRPA1. WT mouse and zebrafish TRPA1 constructs bound CaM in a Ca^2+^-dependent manner (**Fig. 6G**). In contrast, WT *C. elegans* TRPA1 and the partial DCTCaMBE truncations of mouse and zebrafish TRPA1 failed to bind CaM (**Fig. 6G**). These results are consistent with the DCTCaMBE serving as a critical and highly conserved Ca^2+^/CaM binding element in TRPA1 channels.

### Ca^2+^/CaM binding to the TRPA1 DCTCaMBE is critical for rapid channel desensitization

Having discovered the high affinity DCTCaMBE in the TRPA1 C-terminus as well as two mutations (Δ1109-1119 and W1103A) that ablate Ca^2+^/CaM binding to the channel, we next wanted to use these mutations to determine the role Ca^2+^/CaM plays in TRPA1 regulation and how disruption of this interaction confers the hyperactivity observed by Ca^2+^ imaging (**Fig. 2G, H, K**, and **L**). Notably, the W1103A hTRPA1 mutant showed a loss in biotinylation by TurboID-CaM C-lobe demonstrating it does not interact with Ca^2+^/CaM in cells (**Fig. 3D** and **E**, blue). The canonical hTRPA1 Ca^2+^ regulation profile can be observed in *Xenopus laevis* oocytes by two-electrode voltage clamp (TEVC), as previously reported providing a system in which to study our Ca^2+^/CaM binding mutants (**Fig. 7A**)^10,87^. First, we characterized the Δ1109-1119 and W1103A hTRPA1 mutants compared to WT channels for their opening and closing kinetics in the absence of Ca^2+^, their peak current amplitudes, as well their AITC sensitivities to ensure neither mutation compromised channel integrity (**Fig. 7A-D** and **S7**). These control experiments revealed that both the Δ1109-1119 and W1103A hTRPA1 mutants exhibit gating kinetics and peak current amplitudes that were not significantly different from WT hTRPA1 as well as slightly reduced AITC sensitivities (**Fig. 7D**, timepoint i and **S7**). Thus, we compared AITC-evoked WT, Δ1109-1119, and W1103A hTRPA1 activities by TEVC in the presence of extracellular Ca^2+^ to directly discern how loss of Ca^2+^/CaM binding affects channel activity. Application of extracellular Ca^2+^ triggered rapid potentiation of WT, Δ1109-1119, and W1103A hTRPA1 channels with comparable peak current amplitudes, suggesting that loss of Ca^2+^/CaM binding to the TRPA1 DCTCaMBE does not impact potentiation (**Fig. 7A-D**, timepoint ii). While WT hTRPA1 channels subsequently exhibited nearly complete desensitization within 25 seconds of extracellular Ca^2+^ addition (**Fig. 7A**, timepoint iii), hTRPA1^Δ1109-1119^ and W1103A channels remained open with 20-30% or 10-20% their initial activity 5 minutes after extracellular Ca^2+^ application, respectively (**Fig. 7B-D**, timepoint iv). These residual activities could be inhibited with the TRPA1 antagonist A-967079, demonstrating that the ongoing activity was due to hTRPA1 channels (**Fig. 7B-C**, timepoint v). It is possible that Ca^2+^ brought in during prolonged activation of Δ1109-1119 and W1103A hTRPA1 channels triggers delivery of new channels to the plasma membrane or other Ca^2+^-dependent signaling pathways that contribute to the extended activation profile. Thus, we pre-incubated oocytes expressing W1103A hTRPA1 with 500 µM EGTA-AM for 20 minutes before TEVC recordings to buffer global Ca^2+^ concentrations while still allowing for local Ca^2+^ regulation^50,88,89^. These recordings reveal a delayed desensitization profile compared to WT hTRPA1 channels and a faster desensitization profile than W1103A hTRPA1 channels without intracellular Ca^2+^ buffering, demonstrating that prolonged Ca^2+^ entry has additional physiological effects (**Fig. 7E**).

**Figure 7.**
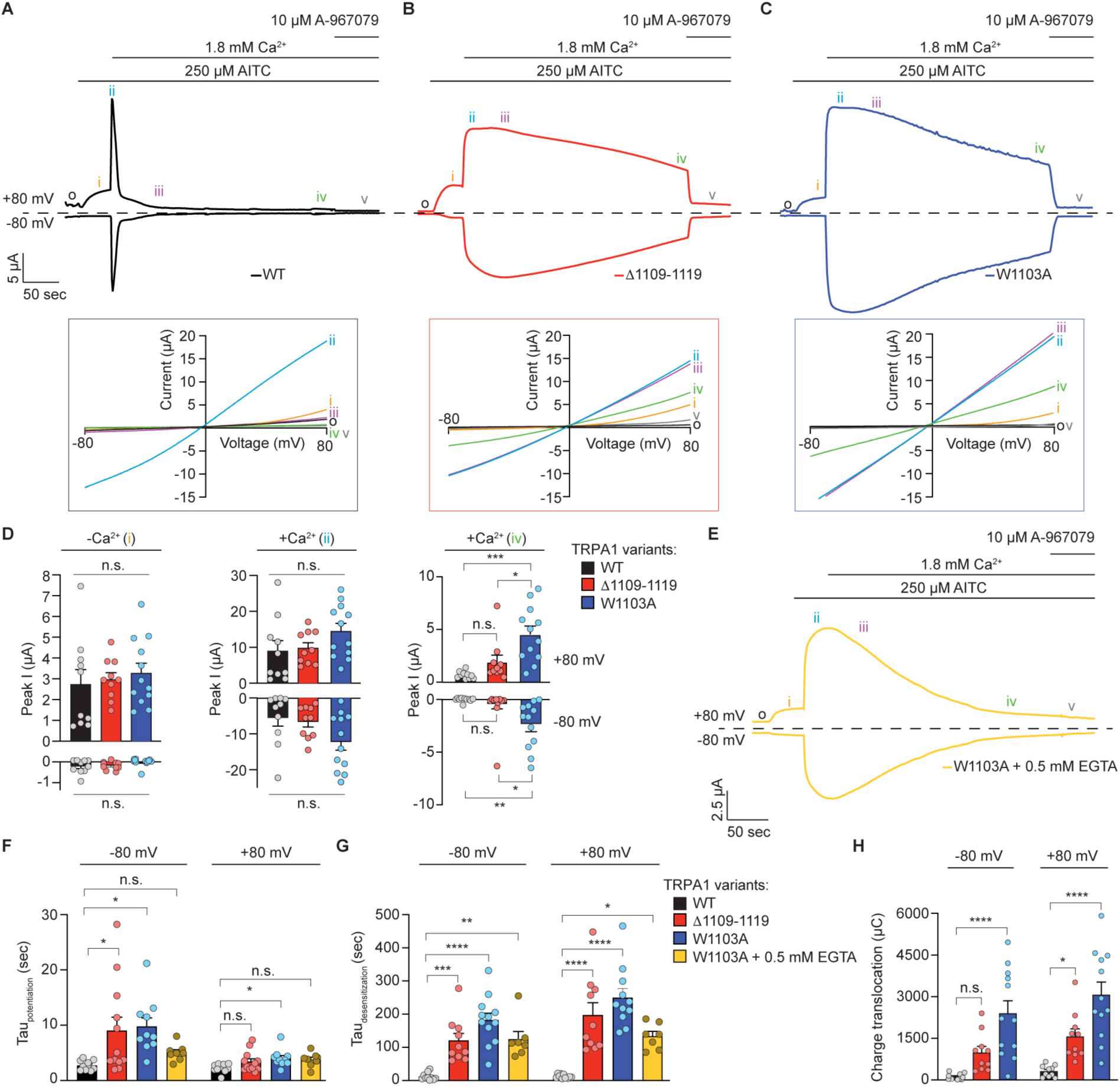
Ca^2+^/CaM binding is critical for TRPA1 desensitization. (**A**-**C**) Representative time-traces at −80 and +80 mV holding potentials from oocytes expressing WT (A, black), Δ1109-1119 (B, red), or W1103A hTRPA1 (C, blue). Current evoked with 250 µM AITC in the absence (orange i) and presence (blue ii) of 1.8 mM extracellular Ca^2+^. Channels were blocked with 10 µM A-967079 (grey v). Dashed line denotes 0 µA current. Protocol of condition application indicated above. Boxed below are the corresponding current-voltage relationships from timepoints indicated by black o (baseline), orange i (AITC without Ca^2+^), blue ii (AITC with Ca^2+^), purple iii (25 seconds after Ca^2+^ addition), green iv (5 minutes after Ca^2+^ addition), and grey v (A-967079 inhibited). (**D**) Quantification of peak current amplitudes at +80 mV (above) and −80 mV (below) before (orange i, left), 25 seconds after (blue ii, middle), and 5 minutes after (green iv, right) Ca^2+^ addition. Colors as indicated in A-C. Data represent mean ± SEM. ***p=0.0002, **p=0.0026, *p=0.0132 or 0.0217 (+80 and −80 mV, respectively), n.s. not significant. (**E**) Representative time-traces at −80 and +80 mV holding potentials from oocytes expressing W1103A hTRPA1 and pre-treated with 0.5 mM EGTA-AM (yellow) 20 minutes prior to recordings to chelate global intracellular Ca^2+^. Dashed line denotes 0 µA current. Protocol of condition application indicated above and timepoints marked as in A-C. (**F** and **G**) Calculated time constants of potentiation (F) and desensitization (G) at −80 mV (left) and +80 mV (right) from fitting data as in A-C and E to a single-exponential function. Data represent mean ± SEM. *p=0.0266 (WT versus Δ1109-1119, −80 mV, tau potentiation), *p=0.0166 and 0.0287 (WT versus W1103A, −80 mV and +80 mV, respectively, tau potentiation), ***p=0.0007 (WT versus Δ1109-1119, −80 mV, tau desensitization), **p=0.0014 (WT versus W1103A with 0.5 mM EGTA, −80 mV, tau desensitization), *p=0.0119(WT versus W1103A with 0.5 mM EGTA, +80 mV, tau desensitization) ****p<0.0001, n.s. not significant. Colors as indicated in legend. (**H**) Quantification of charge translocation (µA•s, µC, *e.g.*, area under the curve) at −80 mV (left) and +80 mV (right) from oocytes used as in A-C. Data represent mean ± SEM. *p=0.0408 (WT versus Δ1109-1119, −80 mV), ****p<0.0001, n.s. not significant (p>0.05). Colors as indicated in legend from panel D. (**D** and **F**-**H**) n= 10 (WT and Δ1109-1119 hTRPA1), 12 (W1103A hTRPA1) or 8 (W1103A hTRPA1 with 0.5 mM EGTA) oocytes per condition, one-way ANOVA with Bonferroni‘s *post hoc* analysis.

To quantitatively characterize the effect of Ca^2+^/CaM binding on proper TRPA1 Ca^2+^ regulation, we calculated rate constants for potentiation and desensitization from baseline-corrected traces at −80 and +80 mV holding potentials. Such calculations reveal that WT hTRPA1 potentiated at a rate of 2.7 sec (−80 mV) and 2.2 sec (+80 mV) and desensitized at a rate of 11.9 sec (−80 mV) and 12.5 sec (+80 mV) following Ca^2+^ addition, consistent with literature precedence (**Fig. 7F** and **G**, black bars)^36,38^. In contrast, Δ1109-1119 and W1103A hTRPA1 exhibited significantly delayed desensitization kinetics with 10- or 15-fold slowing to desensitization with inward currents (*e.g.,* −80 mV) and 16- or 20-fold slowing to desensitization kinetics with outward currents (*e.g.,* +80 mV), respectively, compared to WT hTRPA1 channels (**Fig. 7G**, red and blue bars, respectively). Even when intracellular Ca^2+^ was chelated with 500 µM EGTA, W1103A hTRPA1 exhibited a 10.5-fold and 10.7-fold slowing to desensitization at +80 and −80 mV, respectively compared to WT hTRPA1 channels (**Fig. 7G**, yellow bars). These fold-slowing to Ca^2+^-dependent desensitization kinetics are on the order of that previously reported for a TRPA1 mutant exhibiting 100-fold reduced Ca^2+^ permeability and may, in part, reflect channel rundown independent of true Ca^2+^ desensitization^36^. Potentiation kinetics were more modestly slowed for Δ1109-1119 and W1103A hTRPA1 compared to WT hTRPA1 (∼3-fold each at −80 mV and ∼1.8-fold each at +80 mV), though this can be largely accounted for by the greater effects to desensitization which would inherently provide more time for full potentiation (**Fig. 7F** red and blue bars, respectively). W1103A hTRPA1 with buffered intracellular Ca^2+^ did not exhibit a significant change to potentiation kinetics compared to WT hTRPA1 channels (**Fig. 7F**, yellow bars). Thus, these results suggest that Ca^2+^/CaM binding at the TRPA1 DCTCaMBE specifically affects rapid desensitization.

Such drastic delays to channel desensitization with Δ1109-1119 and W1103A hTRPA1 allows for sustained ion conduction through active mutant channels over a prolonged period, which could have significant physiological effects including delivery of additional channels to the plasma membrane as suggested above. hTRPA1^Δ1109-1119^ channels exhibit a large, albeit nonsignificant increase in translocated charge at −80 mV and a significant, 5-fold increase at +80 mV (**Fig. 7H**, red bars). The W1103A hTRPA1 mutant exhibited significant, 20- and 10-fold increases in translocated charge at −80 and +80 mV, respectively (**Fig. 7H**, blue bars). These collective differences in mutant channel function are not due to changes in total protein expression (**Fig. S1C**).

To independently characterize the effect of CaM binding on hTRPA1 Ca^2+^ regulation, we used the MBP- hTRPA1^1089-1119^ peptide (called the MBP-peptide) as an experimental tool to sequester endogenous CaM since this peptide competed with WT hTRPA1 in CaM binding assays (**Fig. S4C** and **D**). Co-expression of MBP-WT hTRPA1 with the MBP-peptide in HEK293T cells yielded significantly enhanced basal and AITC-evoked activity by Ca^2+^ imaging compared to channels co-expressed with free MBP, presumably due to endogenous CaM sequestration and release from CaM regulation (**Fig. S8A-C**). In contrast, the MBP-peptide had no effect on hTRPA1^Δ1109-1119^ activity consistent with its insensitivity to Ca^2+^/CaM regulation (**Fig. S8D-F**). We next asked how the MBP-peptide affects channel function by injecting oocytes expressing WT hTRPA1 with purified free MBP tag or MBP-peptide at a final concentration of 100 µM one hour prior to TEVC recordings. WT hTRPA1 profiles from oocytes injected with free MBP were indistinguishable from those with WT hTRPA1 alone (compare **Fig. 7A** and **S8G**) and the channels exhibited similar desensitization rates (11.9 and 12.5 sec at −80 and +80 mV for WT hTRPA1, respectively versus 9.1 and 10.34 sec at −80 and +80 mV for WT hTRPA1 with free MBP, respectively). In contrast, WT hTRPA1 channels from oocytes injected with the MBP-peptide exhibited 16- and 18-fold elongations to desensitization kinetics at −80 and +80 mV, respectively, compared to channels from oocytes injected with free MBP (**Fig. S8H-J**). Thus, the MBP-peptide had a comparable effect on hTRPA1 activity as the Δ1109- 1119 and W1103A mutants. Collectively, these results demonstrate that Ca^2+^/CaM is a critical regulator of hTRPA1 wherein Ca^2+^/CaM binding to the DCTCaMBE is required for proper hTRPA1 desensitization in intact cells while having no effect on potentiation. This also provides compelling evidence that potentiation and desensitization are independent regulatory events, as previously proposed^36^.

### High extracellular Ca^2+^ partially rescues desensitization-resistant TRPA1

The free CaM C-lobe has a severely limited ability to respond to Ca^2+^ flux compared to free WT CaM, which raises questions about how it could confer desensitization through binding at the hTRPA1 DCTCaMBE. One possibility is that CaM does not directly transmit a Ca^2+^ signal to TRPA1, but instead its binding primes an intrinsic Ca^2+^ binding site in TRPA1 that triggers channel desensitization. Indeed, TRPA1 can be desensitized by direct perfusion of Ca^2+^ to its cytoplasmic domains in pulled patches, suggesting TRPA1 desensitization involves a direct Ca^2+^ binding event^36,40,90^. If true, desensitization of Ca^2+^/CaM binding deficient TRPA1 mutant channels should be at least partially rescued by increasing the extracellular Ca^2+^ concentration available to permeate through open channels. Consistent with literature precedence, we observed that WT hTRPA1 channels desensitize faster with 10 mM extracellular Ca^2+^ than with 1.8 mM extracellular Ca^2+^ (**Fig. 8A** and **B**)^36,38^. W1103A hTRPA1 channels were even more sensitive to extracellular Ca^2+^ concentration exhibiting an approximately 3-fold faster desensitization rate with 10 mM extracellular Ca^2+^ than with 1.8 mM extracellular Ca^2+^ (**Fig. 8C** and **D**). Despite these enhanced rates, W1103A hTRPA1 channels were still approximately 4-fold slower to desensitize than WT hTRPA1 with 1.8 mM extracellular Ca^2+^. Collectively, these results are consistent with loss of Ca^2+^/CaM binding affecting an intrinsic TRPA1 Ca^2+^ binding site that can only be partially rescued by the higher intracellular Ca^2+^ concentration permeated by W1103A hTRPA1 channels with 10 mM extracellular Ca^2+^. Future work will identify this intrinsic TRPA1 Ca^2+^ binding site and its contribution to channel desensitization.

**Figure 8.**
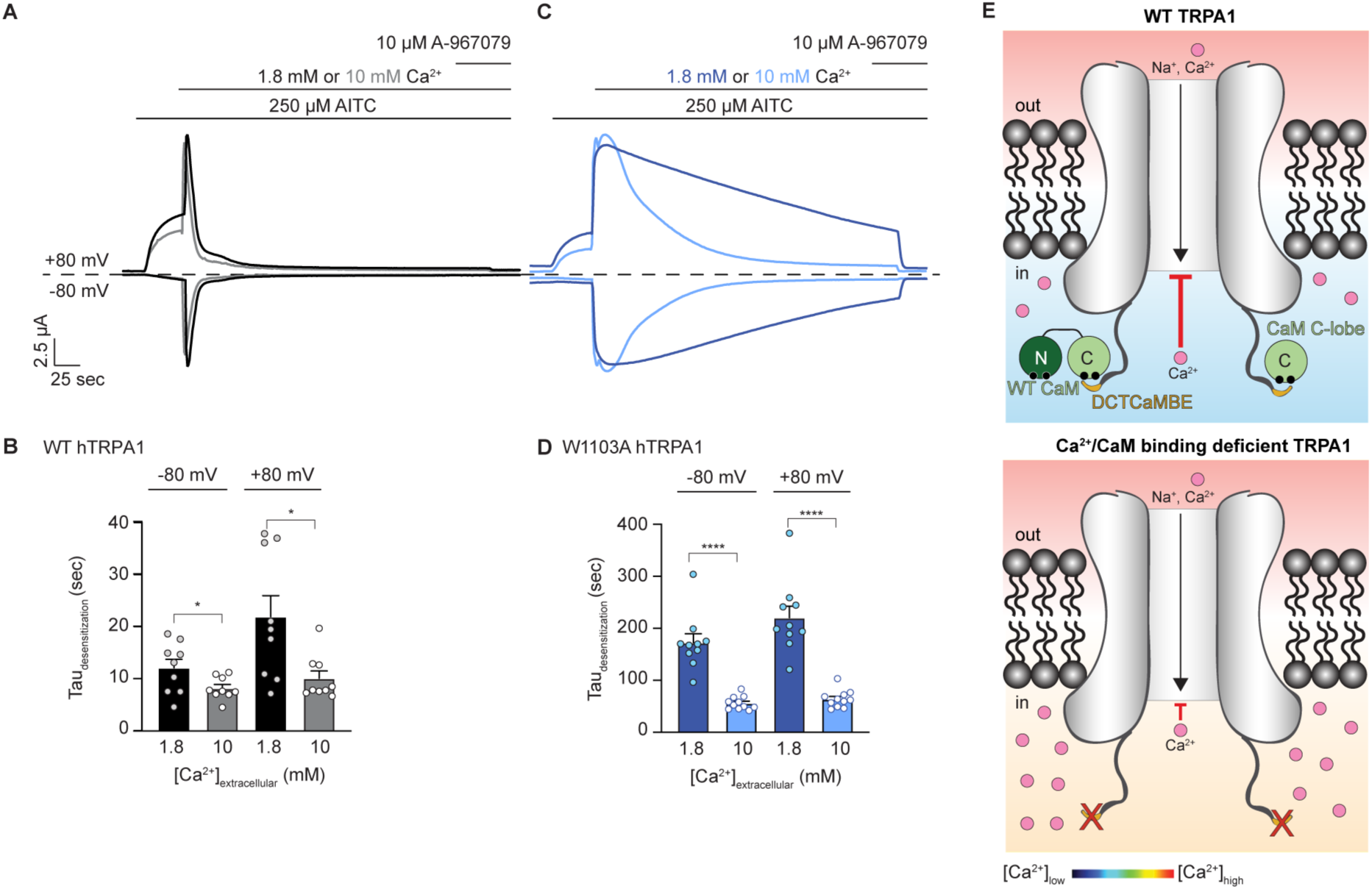
Loss of Ca^2+^/CaM binding at the TRPA1 DCTCaMBE can be partially rescued by high extracellular Ca^2+^. (**A** and **C**) Representative time-traces at −80 and +80 mV holding potentials from oocytes expressing WT (A) or W1103A hTRPA1 (C). Current evoked with 250 µM AITC in the absence and presence of 1.8 mM (black/dark blue) or 10 mM (grey/light blue) extracellular Ca^2+^. Channels were blocked with 10 µM A-967079. Dashed line denotes 0 µA current. Protocol of condition application indicated above. Extracellular solutions contained 125 µM niflumic acid (NFA). (**B** and **D**) Calculated time constants of desensitization from fitting data as in (A) and (C) at 1.8 or 10 mM extracellular calcium to a single-exponential function. Data represent mean ± SEM. ****p<0.001, *p= 0.0424 (WT hTRPA1, −80 mV) or 0.0146 (WT hTRPA1, +80 mV). n= 9 (WT), 10 (W1103A, 1.8 mM Ca^2+^) or 11 (W1103A, 10 mM Ca^2+^) oocytes per condition, two-tailed Student’s t-test. (**E**) Model illustrating Ca^2+^ desensitization of TRPA1 (red inhibition arrow) in a WT channel (top) versus a Ca^2+^/CaM binding deficient channel (bottom). Channel activation facilitates permeation of sodium (Na^+^) and Ca^2+^ ions into the cytoplasm. WT CaM (dumbbell) or the CaM C-lobe (green circle) binding to the TRPA1 DCTCaMBE (yellow) in the distal unstructured C-terminus primes TRPA1 for rapid desensitization following Ca^2+^ permeation (top, blue color on the Ca^2+^ concentration heat map). When the TRPA1:Ca^2+^/CaM interaction is disrupted, channels remain active allowing intracellular Ca^2+^ concentrations to rise (bottom, orange color on the Ca^2+^ concentration heat map) before incomplete desensitization is observed.

## DISCUSSION

TRPA1 is a Ca^2+^-permeable ion channel implicated in chronic pain and inflammation whose activity is controlled by rapid desensitization following a cytosolic Ca^2+^ increase^13,21,23–32,36,39–41^. Despite the importance of desensitization to restricting proper TRPA1 functionality, this regulatory mechanism, and whether it involves the universal Ca^2+^ sensor CaM, has remained controversial^36,39–41,90^. Here, we find that Ca^2+^/CaM binds human TRPA1 best at basal Ca^2+^ concentration, co-localizes with TRPA1 at rest, and suppresses channel activity when co-expressed in heterologous cells, suggesting Ca^2+^/CaM is a *bona fide* TRPA1 regulator. Ca^2+^/CaM mediates this regulation through a previously unidentified, highly conserved, high-affinity binding site in the TRPA1 distal C-terminus that we have named the distal C-terminal CaM binding element (DCTCaMBE). The TRPA1 DCTCaMBE resides in the final 18 amino acids of a 41-amino acid C-terminus that has yet to be resolved by structural studies (*e.g.*, the TRPA1 structurally unresolved C-terminus, **Fig. 1A**)^39,91,92^. Quantitative binding assays, structure predictions, and NMR spectroscopy reveal that the TRPA1 DCTCaMBE exclusively engages the high Ca^2+^-affinity CaM C-lobe in a Ca^2+^- dependent manner, supporting a pre-association of TRPA1 and Ca^2+^/CaM in cells. Mutation of putative TRPA1 DCTCaMBE residues in the full-length channel or sequestration of CaM with a TRPA1 C-terminal peptide ablates Ca^2+^/CaM binding at basal Ca^2+^ concentration and yields hyperactive channels that exhibit a 10- to 20-fold slowing of Ca^2+^-mediated desensitization, which can only be partially rescued by an increased extracellular Ca^2+^ concentration (**Fig. 8E**, bottom). This functional perturbation is more drastic than that caused by all known TRPA1 channelopathic mutations and suggests that disruption of the TRPA1:Ca^2+^/CaM interaction would have profound neurological effects, which could provide a useful tool for studying TRPA1 hyperactivity in genetically tractable animal models^10–12^. Indeed, some TRPV4, Nav1.1, and Nav1.2 channelopathic mutations are thought to mimic loss of CaM binding^93,94^. Our work identifies the TRPA1 DCTCaMBE as the main Ca^2+^/CaM binding element and reveals that Ca^2+^/CaM or Ca^2+^/CaM C- lobe binding to this site is critically required for rapid TRPA1 desensitization (**Fig. 8E**, top).

CaM binds and regulates the activity of hundreds of effector proteins including many ion channels through several types of binding motifs, requiring a remarkable degree of plasticity for a protein that is less than 17 kDa^48,71,76,77,81,84,94–102^. The canonical binding mode involves both lobes of Ca^2+^-bound CaM wrapping around a single effector protein alpha helix through a series of hydrophobic interactions with the CaM lobes’ hydrophobic clefts though the N- and C-lobes can make a bridged interaction by engaging separate effectors or by engaging distinct binding motifs^44,48,71,76,77,81,84,94,97–103^. Our binding studies, AlphaFold predictions, and NMR data support exclusive engagement of the TRPA1 DCTCaMBE by the Ca^2+^/CaM C-lobe. The predicted TRPA1 DCTCaMBE is a 10-residue binding segment within a putative alpha helix with four hydrophobic residues at positions 2, 5, 6 and 9 and basic residues at positions 1 and 10 (**Fig. 6F**). It is predicted to bind the Ca^2+^/CaM C-lobe as an alpha helix with W1103 forming the main hydrophobic anchor akin to Y1494 in the Nav1.5 domain III-IV linker (**Fig. 6B** and **S6F**)^77^. A Dali search reveals that this predicted binding mode is akin to other alpha helical effector peptides bound to the calcified CaM C-lobe (**Fig. S6D**). Following naming conventions, the TRPA1 DCTCaMBE is a 2-5-6-9 motif; it provides another example of CaM binding plasticity and it supports the growing body of literature that non-canonical CaM binding motifs are more common than previously expected^95^.

Our IF imaging and proximity biotinylation data suggest that Ca^2+^/CaM forms a stable interaction with TRPA1 at rest through an interaction between the TRPA1 DCTCaMBE and the CaM C-lobe (**Fig. 3**). These results raise the intriguing possibility that CaM is the first discovered TRPA1 auxiliary subunit. Auxiliary subunits including CaM have been studied extensively and were shown to regulate ion channel folding, stability, trafficking, localization, stimulus sensitivity, and gating kinetics^60–64,104,105^. Unlike KCNQ channels, we did not find that Ca^2+^/CaM affects TRPA1 assembly or trafficking (**Fig. 1G, 1H, 2I** and **2J**)^106–109^. Instead, our work reveals that Ca^2+^/CaM is an indispensable auxiliary subunit required for rapid channel desensitization that effectively tightens the TRPA1 functional window and limits total ion influx (**Fig. 7G** and **8E**). In this way, Ca^2+^/CaM operates akin to the auxiliary subunits of voltage-gated ion channels^47,49–64,110^. Future *in vitro* work on TRPA1 should ensure that CaM is expressed at sufficient levels to initiate rapid desensitization and to avoid artifactual results from hyperactive channels. This is particularly key for heterologous systems where endogenous CaM pools can be easily overwhelmed by overexpressed TRPA1 and must be supplemented by exogenous CaM.

We identified the TRPA1 DCTCaMBE in the distal C-terminus, which has never been resolved structurally, and thus it is unknown where the DCTCaMBE resides in relation to domains that govern channel gating^39,91,92^. In this way, TRPA1 joins the ranks of the many other ion channels with C-terminal CaM regulatory domains^47,49–59,99,110–113^. Unlike these other channels, however, the TRPA1 C-terminus is likely located quite far from the pore. This raises mechanistic questions about how Ca^2+^/CaM binding to the TRPA1 DCTCaMBE could promote channel desensitization (**Fig. 1A** and **B**). One intriguing model is a coordinated response between the TRPA1 DCTCaMBE and the previously identified CaM binding domain (CaMBD)^41^, which resides near the ion conduction pathway, through sequential binding of the CaM C- and N-lobes, respectively as intracellular Ca^2+^ rises to trigger desensitization. Such mechanisms involving disparate binding elements within the same channel for the CaM N- and C-lobes have been proposed for CaM- and Ca^2+^-mediated SK channel activation, TRPV5 and TRPV6 inhibition, as well as regulation of voltage-gated ion channels^42,43,45,46,48,50,77^. However, we and others have shown that the CaM N-lobe is dispensable for TRPA1 regulation (**Fig. 1E** and **F**)^41^.

The lack of dependence of TRPA1 on the CaM N-lobe for rapid desensitization is strikingly distinct. Indeed, most CaM-regulated ion channels rely on conformational changes in CaM upon Ca^2+^ binding to the N- and C-lobes that either directly blocks the pore or induces a conformational change in the channel to regulate gating^42–47,49–59^. Thus, a second possible model is that the Ca^2+^/CaM C-lobe engages the TRPA1 DCTCaMBE with a calcified EF hand 4 and, upon Ca^2+^ influx, Ca^2+^ binding to EF hand 3 triggers a conformational change in the CaM C-lobe that contributes to rapid desensitization. EF hands 3 and 4 were previously shown to both be required for TRPA1 regulation and future mechanistic work will further explore this possible model^41^. A third and intriguing model is that CaM may not be functioning as a direct Ca^2+^ sensor, *per se* in rapid TRPA1 desensitization. Our data show that while ablation of Ca^2+^/CaM binding to the TRPA1 DCTCaMBE drastically slows the desensitization rate, it does not completely ablate desensitization; permeating Ca^2+^ ions are, albeit inefficiently, able to desensitize TRPA1 and this slowed desensitization can be partially rescued by increasing the extracellular Ca^2+^ concentration (**Fig 7** and **8E**, bottom, smaller red inhibition arrow). Collectively, we believe our results support a model wherein Ca^2+^/CaM binding at the DCTCaMBE primes the affinity and/or accessibility of an intrinsic TRPA1 Ca^2+^ binding site so that it can readily respond to Ca^2+^ flux through the channel pore and trigger rapid desensitization (**Fig 8E**, top, larger red inhibition arrow). This model can explain why Ca^2+^ ions perfused directly onto the TRPA1 cytoplasmic domains in excised patches that were previously exposed to EGTA can still trigger desensitization^36,40,90^. Future structural and functional work are needed to identify this Ca^2+^ binding site, to determine how Ca^2+^ binding triggers rapid TRPA1 desensitization, and to uncover how Ca^2+^/CaM affects this process. Finally, while the CaM N-lobe is not required for rapid TRPA1 desensitization, it may serve to enhance Ca^2+^/CaM binding to TRPA1 through the CaMBD or other unidentified site that may contribute to other aspects of channel regulation, or it may remain available to interact with other local effectors and initiate Ca^2+^- and CaM-dependent signaling^114^.

Collectively, we identified a highly conserved and critical Ca^2+^/CaM regulatory element in the human TRPA1 distal unstructured C-terminus that is required for proper channel function. Localization of the TRPA1 DCTCaMBE to the distal C-terminus is serendipitous since it allowed us to completely ablate Ca^2+^/CaM binding through genetic modification of TRPA1 without affecting intrinsic channel function (*e.g.*, WT and mutant hTRPA1 channels exhibited similar activities in Ca^2+^ free solution, **Fig. 7A-D** and **S7**). Thus, we were able to directly reveal this critical role for Ca^2+^/CaM in TRPA1 regulation without relying on perturbation of CaM, which can have off-target effects. Moreover, our work demonstrates how short segments of structurally unresolved amino acids can drastically affect channel function in unexpected ways. Ion channels are highly enriched for intrinsically disordered segments at both their N- and C-termini with some of them, including TRPA1 as described here, shown to be indispensable for proper function ^115,116^. For others, such as the related capsaicin receptor, TRPV1, a structurally unresolved element in the distal C-terminus governs phosphatidylinositol lipid regulation^117,118^ ^119^. Perturbations therein promote TRPV1 sensitization and a splice variant that removes this regulatory element may underlie infrared detection capability in vampire bats^117,119,120^. Given their wide distribution amongst the ion channel proteome including the large TRP channel family, other intrinsically disordered segments are bound to have novel channel regulatory modes^121^. Such segments provide a ripe evolutionary source for new functionality as they are more permissive to mutations than structured regions, which is particularly useful for transmembrane proteins whose complex regulation has developed mainly through extra-membrane domains^87,120,122–124^. While the cryo-EM resolution revolution has been a boon for the ion channel and membrane protein fields, our work illustrates the need to study structurally unresolved and intrinsically disordered regions with renewed interest and vigor.

## METHODS

### Plasmid Construction

DNA sequences for human TRPA1, human TRPA1^448–1078^, and human Calmodulin were PCR amplified and subcloned into a p3xFLAG-eYFP-CMV-7.1 vector (Addgene) at the NotI/BamHI sites using In-Fusion EcoDry cloning (Takara) according to manufacturer protocols. Human TRPA1, mouse TRPA1, zebrafish TRPA1, and C. *elegans* TRPA1 were subcloned into 8xHis-MBP pFastBac1 modified with a CMV promoter (obtained from David Julius’ lab) at the BamHI/NotI sites following the same protocol. 8xHis-MBP pFastBac1 modified with a CMV promoter human TRPA1^1089-1119^ was created by introducing a BamHI site at the −1 position of the codon encoding TRPA1^1089^ and digesting the plasmid with BamHI restriction enzyme followed by gel purification. The linearized vector segment was then transformed into XL-10 Gold cells resulting in a repaired vector. A similar strategy was followed for the creation of 8xHis-MBP pFastBac1 with a CMV promoter human TRPA1^1079-1119^, and human TRPA1^1099-1119^. For bacterial protein purification, DNA sequences for 8xHis-MBP TRPA1^1089-1119^ and 3xFlag-6xHis-WT CaM were PCR amplified and subcloned into a pET28a vector (Addgene) at the NcoI/NotI sites using In-Fusion EcoDry cloning. All mutants including hTRPA1^Δ1109-1119^, hTRPA1^Δ1079-1119^, hTRPA1^Δ1089-1119^, hTRPA1^Δ1099-1119^, and hTRPA1^Δ1-61^, TRPA1^W1103A^, TRPA1^L1107A^, TRPA1^V1110A^, TRPA1^Quad^, TRPA1^RK/AA^, CaM_12_, CaM_34_, and CaM_1234_ and CaM C-lobe were created using QuikChange Lightning site-directed mutagenesis (Agilent). 3xFLAG-TurboID-CaM C-lobe was generated by producing a MegaPrimer for TurboID that would replace the CaM N-lobe following a site-directed mutagenesis reaction with the 3xFLAG-WT hCaM template. 3xFLAG-TurboID was generated from 3xFLAG-TurboID-CaM C-lobe by site-directed mutagenesis to replace the CaM C-lobe sequence with a TAG stop codon. V5-tagged CaM was created by site-directed mutagenesis to replace the 3xFLAG-tag in the construct above. For expression in *Xenopus laevis* oocytes, 3xFLAG-hTRPA1 variant genes were subcloned into the combined mammalian/oocyte expression vector pMO (obtained from David Julius’ lab) prior to generating cRNAs.

All DNA primers were ordered from ThermoFisher and all constructs were sequence-verified using the Yale School of Medicine Keck DNA Sequencing Facility.

### Mammalian Cell Culture and Protein Expression

Human embryonic kidney cells (HEK293T, ATCC CRL-3216) were cultured in Dulbecco’s modified Eagle’s medium (DMEM; Invitrogen) supplemented with 10% calf serum and 1x Penicillin-Streptomycin (Invitrogen) at 37°C and 5% CO_2_. Neuro2A cells (ATCC CCL-131) were cultured in Dulbecco’s modified Eagle’s medium (DMEM; Invitrogen) supplemented with 10% fetal bovine serum and 1x Penicillin-Streptomycin (Invitrogen) at 37°C and 5% CO2. Cells were grown to ∼85-95% confluence before splitting for experiments or propagation. HEK293T cells cultured to ∼95% confluence were seeded at 1:10 or 1:20 dilutions into 6- or 12-well plates (Corning), respectively. After 1-5 hours recovery, cells were transiently transfected with 1 μg plasmid using jetPRIME (Polyplus) according to manufacturer protocols. For surface biotinylation experiments, cells were co-transfected with 0.5 µg 3xFLAG-tagged hTRPA1 variants and 0.5 µg 3xFLAG- CaM variants.

### Cell Lysis and Pulldown Experiments

16-24 hours post-transfection, HEK293T cells were washed with PBS (calcium and magnesium free) and lysed in 75-150 μL of TRPA1 lysis buffer (40 mM Tris pH 8.0, 150 mM NaCl, 5 mM DDM, 500 μM EGTA, EDTA-free cOmplete protease cocktail inhibitor tablet) at 4°C while gently nutating. Cell debris were pelleted from the resulting lysates by centrifugation at 15,000 RPM for 10 minutes at 4°C. Total protein concentration in lysates were quantified using a BCA assay (Pierce). Equal concentrations of protein lysate (100 μg) from each experimental condition were added to resins as specified below. 10% of loaded protein amount was reserved as a whole-cell lysate loading control. Buffers were supplemented with free Ca^2+^ as indicated in each figure. Concentrations of CaCl_2_ required for each free Ca^2+^ concentration were calculated using Ca-EGTA calculator v1.3 using constants from Theo Schoenmakers’ Chelator on the MaxChelator website^125^.

### CaM-Agarose Pulldown

100 µg lysates were incubated with 10 μL of lysis buffer-equilibrated Calmodulin-agarose (Sigma-Aldrich) for 1 hour at 4°C with gentle nutation. Resin beads were washed 3-5 times with lysis buffer prior to elution; resin was pelleted, and bound proteins were eluted in TRPA1 lysis buffer supplemented with 5 mM EGTA for 30 minutes at 4°C with gentle nutation.

### Biotinylation Assays

Surface biotinylation assays were conducted as previously reported^10^. Briefly, HEK293T cells were seeded in a 6-well plate pre-coated with Poly-L-Lysine (Sigma) and transfected with expression plasmids. 16-24 hours post-transfection, cells were washed with PBS, placed on ice, and incubated for 20 minutes with chilled 0.05 mg/mL Sulfo-NHS-LC-Biotin (ThermoFisher) in Ringer’s solution. Cells were then washed with chilled wash buffer (PBS + 100 mM Glycine) and the reaction was quenched on ice for 30 minutes with 100 mM Glycine and 0.5% BSA in PBS. Cells were then washed three times and lysed in TRPA1 lysis buffer containing 100 mM Glycine directly on the 6-well plate. Lysates were collected, protein concentration was determined by BCA assay, biotinylated proteins were isolated by Neutravidin pulldown and analyzed by immunoblot as described below. During quantification, any FLAG signal from CaM in the eluates were subtracted from all other conditions to account for probe internalization.

For proximity biotinylation assays, HEK293T cells co-expressing 3xFLAG-TurboID or 3xFLAG-TurboID- CaM C-lobe and 3x-FLAG-tagged WT, Δ1109-1119, or W1103A hTRPA1 were treated with 50 uM biotin supplemented into culture media ∼48 hours post-transfection. Cells were lysed 15 minutes later with TRPA1 lysis buffer and affinity purified *via* the 3xFLAG tags as described below. Biotinylated proteins were detected with HRP-conjugated Streptavidin (GE, 1:50,000).

### Neutravidin Pulldown

Cell lysates that were generated following surface labeling experiments were incubated with 15 μL of lysis buffer-equilibrated Neutravidin resin (Pierce) for 2 hours at 4°C with gentle nutation. The resin was then washed with lysis buffer three times, followed by a harsher wash with 1x PBS + 100 mM DTT. Resin was then washed once each with lysis buffer and 1x PBS. Biotinylated protein was eluted from the resin with a multi-step protocol to prevent TRPA1 aggregation while maximizing protein elution from the resin. First, resin was incubated with 10 μL of biotin elution buffer (TRPA1 lysis buffer, 100 mM Glycine, 10 mM Biotin, 1% SDS) on ice for 10 min, followed by addition of 1 μL ß-mercaptoethanol (BME; Sigma) to each sample and incubation on ice for 5 minutes, and finally by addition of 4 μL 4x Laemmli buffer + 10% BME with incubation at 65°C for 10 minutes. The resin was centrifuged, supernatant was removed and combined with additional 4 μL Laemmli buffer + 10% BME for SDS-PAGE analysis.

### FLAG Immunoprecipitation

Lysates were incubated with EZview Red Anti-FLAG M2 affinity resin (Sigma) for 1 hour at 4°C with gentle rotation. Resin beads were washed four times with TRPA1 lysis buffer prior to elution with 125 µg/ml 3xFLAG peptide (Sigma).

### SDS-PAGE and Immunoblot

Samples were combined with Laemmli Sample buffer + 10% BME and separated on pre-cast 4-20% SDS-PAGE gels (Bio-Rad). Gels were transferred onto PVDF membranes (Bio-Rad) by semi-dry transfer at 15V for 40 minutes. Blots were blocked in 3% BSA prior to antibody probing. The following primary antibodies were used in PBST buffer (Boston Bioproducts): anti-MBP (mouse, 1:30,000, New England Biolabs), anti-FLAG (mouse, 1:30,000, Sigma), anti-tubulin (mouse, 1:5,000 in 3% BSA, Sigma). HRP-conjugated IgG secondary anti-mouse antibody was used as needed (rabbit, 1:25,000, Invitrogen). Membranes were developed using Clarity Western ECL substrate (Bio-Rad) and imaged using a Chemidoc Imaging System (Bio-Rad). Densitometric quantifications were performed with ImageJ software. All quantified band intensities for eluted samples were divided by their tubulin-normalized input band intensities.

### Immunofluorescence Imaging and Co-Localization Analysis

Neuro2A cells (ATCC CCL-131) transiently transfected with jetPRIME (Polyplus) according to manufacturer protocols were transferred to poly-L-lysine-coated cover slips and incubated for 16-20 hours prior to immunostaining. Cells were fixed on coverslips with 4% paraformaldehyde for 20 minutes, then permeabilized with 0.1% Triton in PBS for 10 min. Cells were then blocked in 4% BSA in PBS followed by incubation in primary anti-V5 monoclonal antibody (mouse, 1:500, Invitrogen) in 4% BSA in PBS at room temperature for 45 min. Cells were then washed with PBS and incubated with Alexa-Fluor488-conjugated secondary anti-mouse antibody in 4% BSA in PBS at room temperature for 45 min (goat, 1:700, Invitrogen), followed by PBS washes. Finally, cells were incubated in Cy3-conjugated anti-FLAG monoclonal antibody (mouse, 1:500, Sigma) in 4% BSA in PBS at room temperature for 45 min. Images were acquired on a Zeiss Axio Observer Z1 inverted microscope equipped with an Airyscan detection unit, using a Plan-Apochromat 63x/1.40 NA oil objective. Image deconvolution was performed with ZEN 2.1 software. Pixel intensity-based co-localization analysis was performed using the colocalization module in ZEN, from which Pearson’s correlation coefficients were calculated for each analyzed cell. Co-localization scatterplot crosshairs were set based on mono-stained samples for green and red channels.

### Ratiometric Calcium Imaging

16-24 hours post-transfection, HEK293T cells were plated into isolated silicone wells (Sigma) on poly-L-lysine (Sigma)-coated cover glass (ThermoFisher). Remaining cells were lysed for anti-FLAG immunoblotting to ensure equivalent expression. 1 hour later, cells were loaded with 10 μg/mL Fura 2-AM (ION Biosciences) in Ringer’s solution (Boston Bioproducts) with 0.025% Pluronic F-127 (Sigma) and incubated for 1 hour at room temperature, then rinsed twice with Ringer’s solution. Ratiometric calcium imaging was performed using a Zeiss Axio Observer 7 inverted microscope with a Hamamatsu Flash sCMOS camera at 20x objective. Dual images (340 and 380 nm excitation, 510 nm emission) were collected and pseudocolour ratiometric images were monitored during the experiment (MetaFluor software). After stimulation with agonist, cells were observed for 45-100 s. AITC was purchased from Sigma and was freshly prepared as a stock at 4x the desired concentration in 1% DMSO and Ringer’s solution. 5 μL of 4x agonist was added to wells containing 15 μL Ringer’s solution to give the final 1x desired concentration. For all experiments, a minimum of 60-90 cells were selected per condition per replicate for ratiometric fluorescence quantification in MetaFluor with 3-5 replicates per experiment. Background signal was quantified from un-transfected cells and subtracted from quantified cells for normalization.

### Bacterial Protein Expression and Purification

Bacterial expression vectors were transformed into BL21(DE3) cells and plated on Kanamycin (Kan) LB agar plates. Individual colonies were selected to inoculate LB Kan overnight starter cultures from which 10 mL was used to inoculate 1 L of Terrific Broth (TB) Kan. Once the inoculated TB Kan reached an OD of 0.8-1.0, protein expression was induced using 1 mM IPTG (GoldBio). After 4 hours post-induction, bacteria were harvested at 3,000 rpm for 20 min, the supernatant was discarded, and cell pellets were snap frozen with liquid nitrogen and stored at −80°C. Bacteria were cultured at 37°C and 220 RPM for each step.

Bacterial pellets were thawed at 4°C and resuspended in bacterial lysis buffer (50 mM Tris pH 8.0, 500 mM NaCl, 2 mM CaCl_2_, 5 mM β-ME, 1 mM PMSF, 10% glycerol, and 5 mg bovine DNase I). Bacteria were lysed on ice using a FisherBrand Model 120 Sonic Dismembrator (Fisher Scientific). The sonicator was set to 50% amplitude and cycled between 1 minute on and 1 minute off for a total of 20 minutes. Bacterial debris were pelleted using an Eppendorf centrifuge 5810 R that had been cooled to 4°C at 3900 RPM for 30 minutes. All constructs contained either a 6xHis- or 8xHis-tag and were purified by gravity-flow immobilized metal affinity chromatography (IMAC) using lysis buffer-equilibrated HisPur Ni-NTA resin (ThermoFisher). To remove non-specifically bound proteins, the resin was washed with low-molar imidazole wash buffer (50 mM Tris pH 8.0, 150 mM NaCl, 5 mM β-ME, 0.1 mM PMSF, and 20 mM imidazole). Bound protein was eluted in 2 mL fractions with high-molar imidazole elution buffer (50 mM Tris pH 8,0, 150 mM NaCl, 2 mM CaCl_2_, 5 mM β-ME, 0.1 mM PMSF, and 300 mM imidazole) and an A_280_ measurement was taken for each fraction using a NanoDrop One (ThermoFisher). Protein-containing fractions were pooled, exchanged into storage buffer (50 mM Tris pH 8.0 150 mM NaCl, 2 mM CaCl_2_, and 0.1 mM PMSF) or oocyte injection buffer (50 mM Tris pH 8.0, 150 mM KCl, and 0.1 mM PMSF) by dialysis overnight or diafiltration and concentrated using Amicon Ultra centrifugal filter with a 10,000 Da molecular weight cutoff (Millipore). Diafiltration is a technique where macromolecules are concentrated using a spin filter and subsequently diluted with the new buffer. This technique is repeated several times and allows for rapid buffer exchanges relative to dialysis methods. CaM and CaM_12_ were further purified through size exclusion chromatography using a Superdex 75 Increase 10/300 GL (Cytiva). MBP-tagged TRPA1^1089-1119^ was incubated with TEV protease overnight at 4°C to cleave the MBP tag. After TEV protease digestion, TRPA1^1089-1119^ was isolated from free MBP using an Amicon Ultra centrifugal filter with a 30,000 Da molecular weight cutoff. SDS-Page and Coomassie staining was used for post-hoc analysis of each step of the purifications.

### Size-Exclusion Chromatography Binding Assay

Each condition was prepared using 100 μM of each purified protein in a total volume of 500 μL using storage buffer as described above and mixed by gentle nutation at 4°C for 5 minutes. Samples were then loaded onto a Superdex 75 Increase 10/300 GL column pre-equilibrated with either storage buffer or storage buffer supplemented with 5 mM EGTA and run at a flow rate of 0.5 mL/min. The A_280_ signal was monitored by the ÄKTA pure chromatography system (Cytiva), and fractions corresponding to observed peaks were collected, concentrated with the appropriate Amicon Ultra cut-off spin filter, and subjected to Coomassie stain analysis.

### Isothermal Titration Calorimetry

All ITC experiments were carried out using a MicroCal VP-ITC (Malvern) at the Keck Biophysical Resource Center at Yale. The cell was loaded with 15 μM CaM or CaM_12_ and the titrant syringe with 810 μM hTRPA1^1089-1119^. All proteins were buffer matched with storage buffer as described above. 35 injections of 3 μL were performed with 240 seconds between each injection and a stir speed of 290 RPM. Data were analyzed with Origin7 software and fit using the one-site model.

### Structure Prediction

We used AlphaFold2-multimer with default parameters and Amber relaxation to generate all models^82,83^. Computation was performed on the Farnam cluster at Yale Center for Research Computing. CaM amino acids 1 – 79 and 80 – 149 were used as the sequences for the calmodulin N- and C-lobe, respectively. For human TRPA1, aa. 993 – 1010, aa. 977 – 1010, and aa. 1092 – 1119 were used as the sequences for the CaMBD, TRP-CaMBD, and DCTCaMBE respectively. Sequences for Human CaM and TRPA1 were retrieved from Uniprot with accession code P0DP23 and O75762. After obtaining predictions from AlphaFold, we used the model with the highest confidence, as judged by average pLDDT, for all further analysis.

### NMR Spectroscopy

NMR spectra were recorded at 25 °C on a Varian Inova 700 MHz spectrometer equipped with a triple resonance cryoprobe. ^15^N-labeled samples were expressed and purified as described above except that cells were grown in M9 medium with the addition of ^15^N-labeled ammonium chloride. The purified samples were exchanged into the storage buffer supplemented with 10% (v/v) D_2_O. Concentration of the samples ranged from 200-700 µM. Data were collected in 3- or 5-mm tubes for samples with concentration lower than 300 or higher than 400 µM, respectively. Binding between the ^15^N-labeled CaM variants and unlabeled hTRPA1^1089-1119^ peptide was monitored by two-dimensional ^15^N−^1^H HSQC experiments. The number of scans and time domain points of the^15^N−^1^H HSQC experiments on samples with concentration greater than 300 µM were set to 8 and 64, respectively. For samples lower than 300 µM, the number of scans and time domain points were set to 64 and 256, respectively. Chemical shift perturbations (CSPs) of the amide protons and nitrogen was monitored to determine binding. CSP was calculated using the formula:

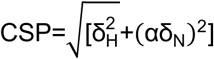

where H and N represents ^1^H and ^15^N nuclei, respectively

*α* is the scaling factor of ^15^N ppm (set to 0.14)

*δ* is the change in chemical shift of the respective nuclei

The spectra were processed in NMRFx analyst^126^.

### Oocyte Electrophysiology

Experiments were conducted as previously reported^10^. pMO vectors carrying 3×FLAG-tagged hTRPA1 constructs were linearized with PmeI, cRNAs were generated by *in vitro* transcription with the mMessage mMachine T7 transcription kit (Thermo) according to the manufacturer’s protocol and were purified with a RNeasy kit (Qiagen). cRNA transcripts were microinjected into surgically extracted *Xenopus laevis* oocytes (Ecocyte) with a Nanoject III (Harvard Apparatus). Oocytes were injected with 0.25 ng (WT hTRPA1) or 0.1 ng (DCTCaMBE variants) of cRNA per cell, and whole-cell currents were measured 24h post-injection using two-electrode voltage clamp (TEVC). Currents were measured using an OC-725D amplifier (Warner Instruments) delivering a ramp protocol from −100 mV to +100 mV applied every second. Microelectrodes were pulled from borosilicate glass capillary tubes and filled with 3 M KCl. Microelectrode resistances of 0.7–1.2 MΩ were used for all experiments. Bath solution contained (in mM) 93.5 NaCl, 2 KCl, 2 MgCl_2_, 0.1 BaCl_2_, and 5 HEPES (pH 7.5). For experiments in the presence of calcium, BaCl_2_ was replaced with 1.8 mM CaCl_2_. For experiments comparing 1.8 and 10 mM extracellular calcium, BaCl_2_ was replaced with 1.8 or 10 mM CaCl_2_ and supplemented with 125 µM niflumic acid (NFA) to inhibit endogenous calcium-activated chloride channels. For global calcium chelation experiments oocytes were incubated with 0.5 mM EGTA-AM for 20 minutes in calcium-free buffer before recordings were taken. Opening and closing times for hTRPA1 constructs were obtained by applying 100 µM AITC in calcium-free buffer to the oocytes for 10 seconds followed by AITC washout with calcium-free buffer. For experiments in Fig. S8, the purified free MBP tag or MBP-hTRPA1^1089-1119^ peptide in oocyte injection buffer were microinjected at least 1 hour prior to recording with a final concentration of 100 μM assuming an oocyte volume of 1 μL. Data were subsequently analyzed using pClamp11 software (Molecular Devices). Oocytes were individually collected after recordings, lysed in 20 µL TRPA1 lysis buffer, and subjected to anti-FLAG immunoblot analysis to confirm construct expression.

### Sequence Alignment

The sequence alignment in Figure 6F was built by aligning the mouse, rat, chicken, fruit bat, ball python snake, boa snake, rat snake, and *Drosophila melanogaster* TRPA1 C-terminal sequences to residues 1089- 1112 of human TRPA1 in Sequence Logo.

### Statistical Analysis

All data quantification was performed in Microsoft Excel. Quantified data presentation and statistical analyses were performed in GraphPad Prism. Criterion for statistical significance for all tests was p<0.05.

## Data Availability

The data that support the findings of this study are available from the corresponding author upon reasonable request. The 6V9W (https://www.rcsb.org/structure/6v9w), 6X2J (https://www.rcsb.org/structure/6x2j), and 4DJC (https://www.rcsb.org/structure/4DJC) PDB files were used in this study. Models were built with ChimeraX version 1.7.1.

## Acknowledgements

We thank Wendy Gilbert, Franziska Bleichert, Lily Kabeche, Tony Koleske, and Yong Xiong for constructive suggestions, Fred Sigworth, and members of the Paulsen lab for helpful discussions and for critical reading of the manuscript, and Grover Paulsen-Sharpe and Marilee Pawsen for moral support. We thank Ewa Folta-Stogniew and Eric Paulson for technical assistance with ITC experiments and NMR data collection, respectively. J.H.S. is supported by an NIH NINDS NRSA pre-doctoral fellowship (1F31NS122412) and a Cellular and Molecular Biology pre-doctoral training grant (T32GM007223). K.M.T. is supported by an Anderson Postdoctoral Fellowship from Yale University. A.B. is supported by a Biophysics pre-doctoral training grant (5T32GM008283-33). This research was supported by an International Association for the Study of Pain Early Career Research Grant, a Rita Allen Foundation and American Pain Society Pain Scholar Award, and by NIH grant R35GM142825 to C.E.P. The content is solely the responsibility of the authors and does not necessarily represent the official views of the National Institutes of Health.

## Author Contributions

J.H.S. and C.E.P. planned the project, designed experiments, cloned all constructs used in the study, and carried out calcium imaging. J.H.S., G.A.A., and C.E.P. carried out Calmodulin binding assays. J.H.S. carried out surface biotinylation, size exclusion chromatography assays, isothermal titration calorimetry, and two-electrode voltage clamp work. J.H.S. and K.M.T. performed protein purifications. A.B. carried out immunostainings. A.B., K.M.T., and C.E.P. carried out proximity biotinylation assays. K.M.T. collected and analyzed ^15^N-^1^H HSQC NMR spectroscopy data. K.M.T. and Y.Z. ran the AlphaFold2 Multimer predictions. J.H.S. and C.E.P. wrote the manuscript with input from K.M.T., G.A.A., A.B., and Y.Z.

