## Supplemental Figures for "Calmodulin binding is required for calcium mediated TRPA1 desensitization"

---

• Supplementary Figures and Figure Legends

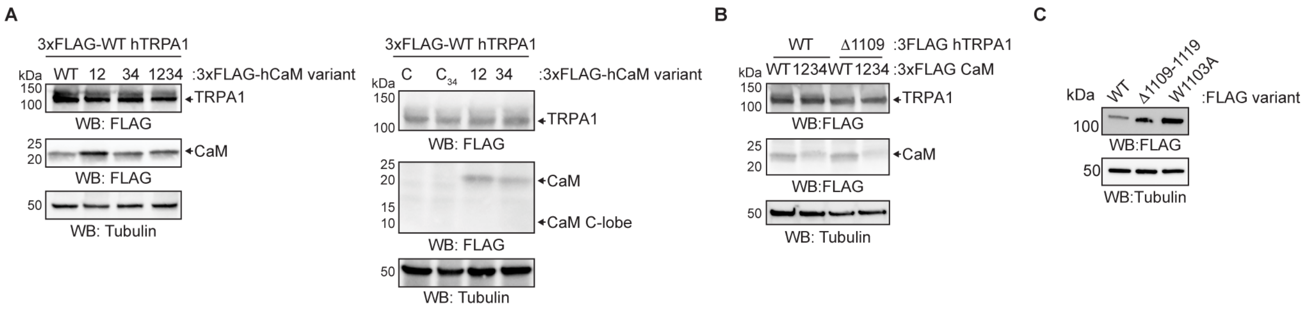


**Figure S1.** Expression tests from functional assays in Fig. 1E (**A**), Fig. 2K (**B**), and Fig. 7 (**C**). Immunoblotting analysis of indicated 3xFLAG-tagged hTRPA1 or hCaM constructs expressed in HEK293T cells (A and B) or *Xenopus laevis* oocytes (C). Samples were probed using an HRP-conjugated anti-FLAG antibody. Tubulin was the loading control.

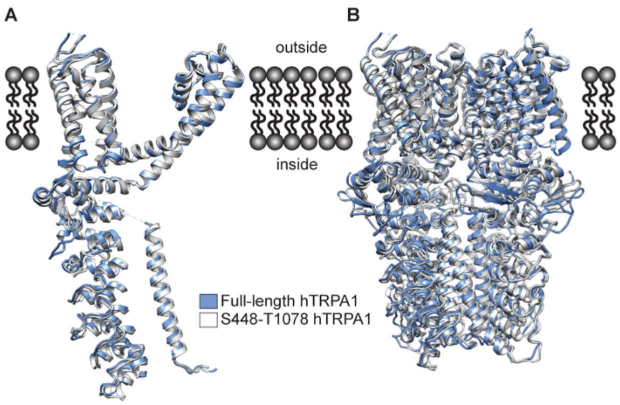


C

| PDB Accession Code | RMSD (Å) | Description of Structure |
|---|---|---|
| 6V9Y | 1.0 | A-967079 |
| 6PQQ | 1.0 | C621S in apo state |
| 6V9W | 1.3 | Ligand-free with bound $Ca^{2+}$ |
| 3J9P | 1.3 | AITC-treated, closed state |
| 6PQP | 1.3 | Covalent BITC |
| 6PQO | 1.4 | Covalent JT010 |
| 6V9V | 1.5 | Covalent Bodipy-iodoacetamide with bound $Ca^{2+}$ |
| 6V9X | 2.8 | Covalent Bodipy-iodoacetamide with open pore |

**Figure S2.** Structural conservation of WT hTRPA1 and minimal $hTRPA1^{448\text{-}1078}$. (**A** and **B**) Ribbon diagram of WT hTRPA1 monomeric (A) or tetrameric (B) atomic models for residues S448-T1078 (blue) overlaid with minimal $hTRPA1^{448\text{-}1078}$ construct (white). Models built with the human hTRPA1 Cryo-EM structure (PDB: 6V9W) or the minimal $hTRPA1^{448\text{-}1078}$ construct (PDB: 6X2J) in ChimeraX. Models were aligned in real space. (**C**) RMSD of $TRPA1^{488\text{-}1078}$ against all available full-length TRPA1 structures. Calculations were performed by the Dali server using $TRPA1^{488\text{-}1078}$ as the search model in a heuristic PDB search.

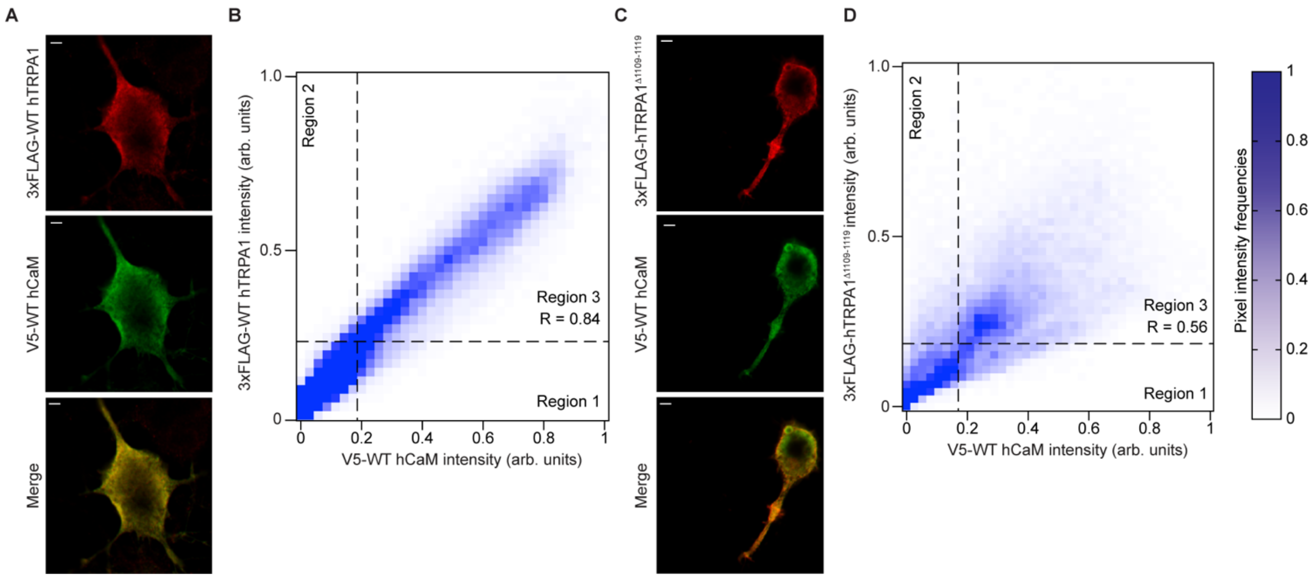


**Figure S3**. TRPA1 and CaM co-localize in cells. (**A** and **C**) Representative raw Airyscan images of Neuro2A cells co-expressing 3xFLAG-WT hTRPA1 (A) or hTRPA1$^{\Delta 1109\text{-}1119}$ (C) with V5-WT CaM as in Fig. 3A. Cells were stained with anti-FLAG (red) and anti-V5 (green) antibodies. Scale bar indicates 2 µm. Images are representative of 3 independent experiments. (**B** and **D**) Heatmaps of 3xFLAG-WT hTRPA1 (B) or hTRPA1$^{\Delta 1109\text{-}1119}$ (D) and V5-WT CaM fluorescence intensity in the Neuro2A cells shown in A and C. Pearson's correlation coefficients (R) were determined using raw images as in A and C and were calculated from Region 3, which exhibited both red and green pixel intensities above background (determined from Regions 1 and 2). Cells are representative of 40 quantified across 3 independent experiments. n > 10,000 pixels in 1 cell per condition.

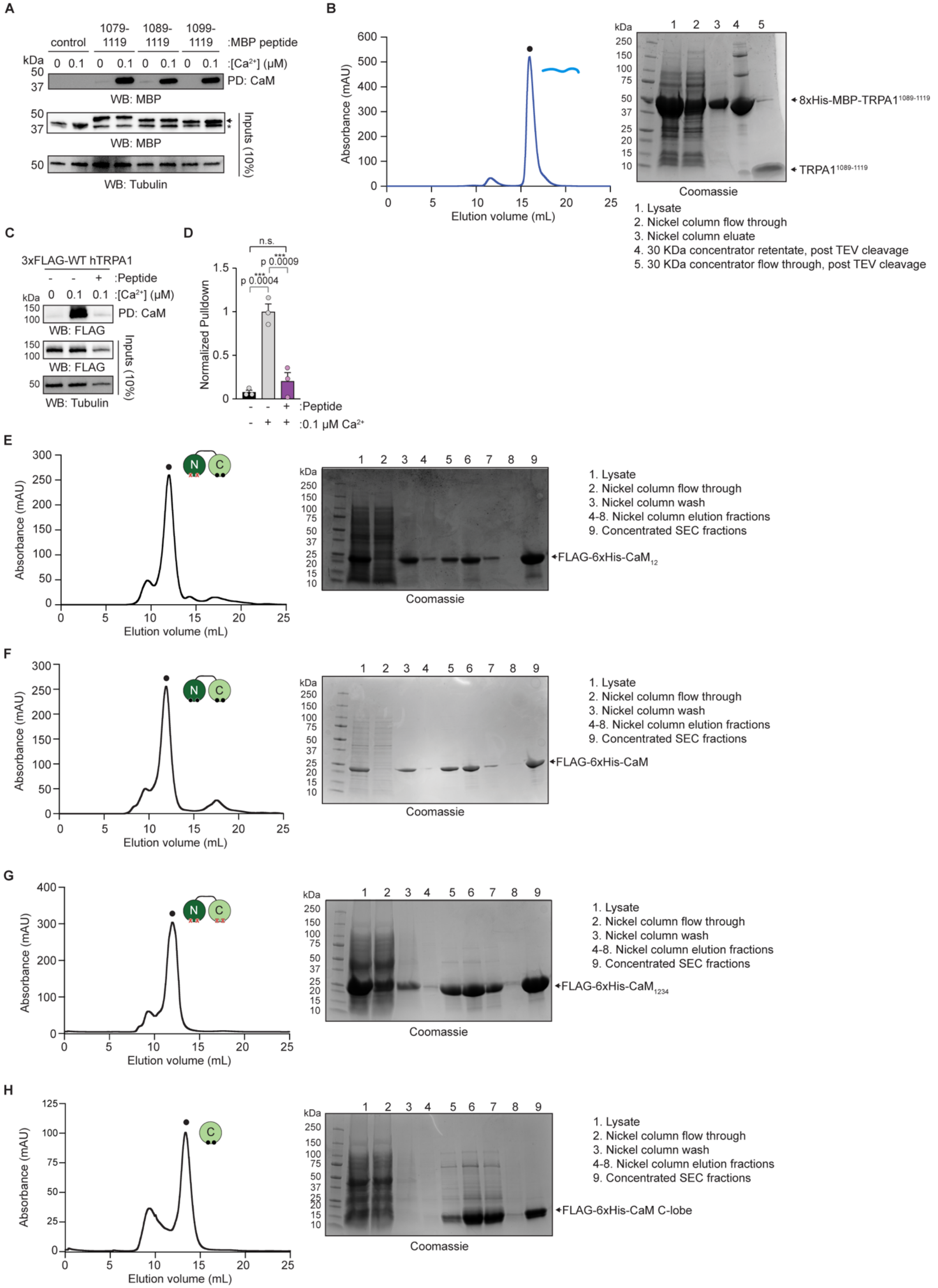


**Figure S4.** (**A**) Immunoblotting analysis of the indicated MBP-tagged hTRPA1 peptide constructs after CaM-agarose pulldown in the absence or presence of $Ca^{2+}$ from lysates of HEK293T cells transfected with free MBP (control), MBP-tagged $hTRPA1^{1079\text{-}1119}$, $hTRPA1^{1089\text{-}1119}$, or $hTRPA1^{1099\text{-}1119}$ peptides. Samples were probed using an anti-MBP primary antibody and an HRP-conjugated anti-mouse secondary antibody. Tubulin from whole cell lysates (10%, inputs) was the loading control. Asterisk (*) denotes free MBP. Arrow denotes MBP-tagged peptide. Purifications of (**B**) the hTRPA1$^{1089-1119}$ peptide, (**E**) WT CaM, (**F**) $CaM_{12}$, (**G**) $CaM_{1234}$, and (**H**) CaM C-lobe for size exclusion chromatography and isothermal titration calorimetry. Representative Superdex 75 chromatograms (left) and Coomassie gels (right) from a purification for each. The peak corresponding to each protein is indicated (black dot). (**C**) Immunoblotting analysis of 3xFLAG-WT hTRPA1 after CaM-agarose pulldown at the indicated $Ca^{2+}$ concentrations in presence or absence of 10 µM hTRPA1$^{1089-1119}$ peptide. Samples were probed as in Fig. 1C. Blot is representative of three independent experiments. (**D**) Quantification of CaM-agarose pulldowns represented in (C). Pulldown was normalized to the WT hTRPA1 with $Ca^{2+}$ and without hTRPA1$^{1089-1119}$ peptide average. Data represent mean ± SEM. ***$p<0.001$, *$p<0.05$, n.s. not significant. n = 3 independent experiments, one-way ANOVA with Tukey's *post hoc* analysis.

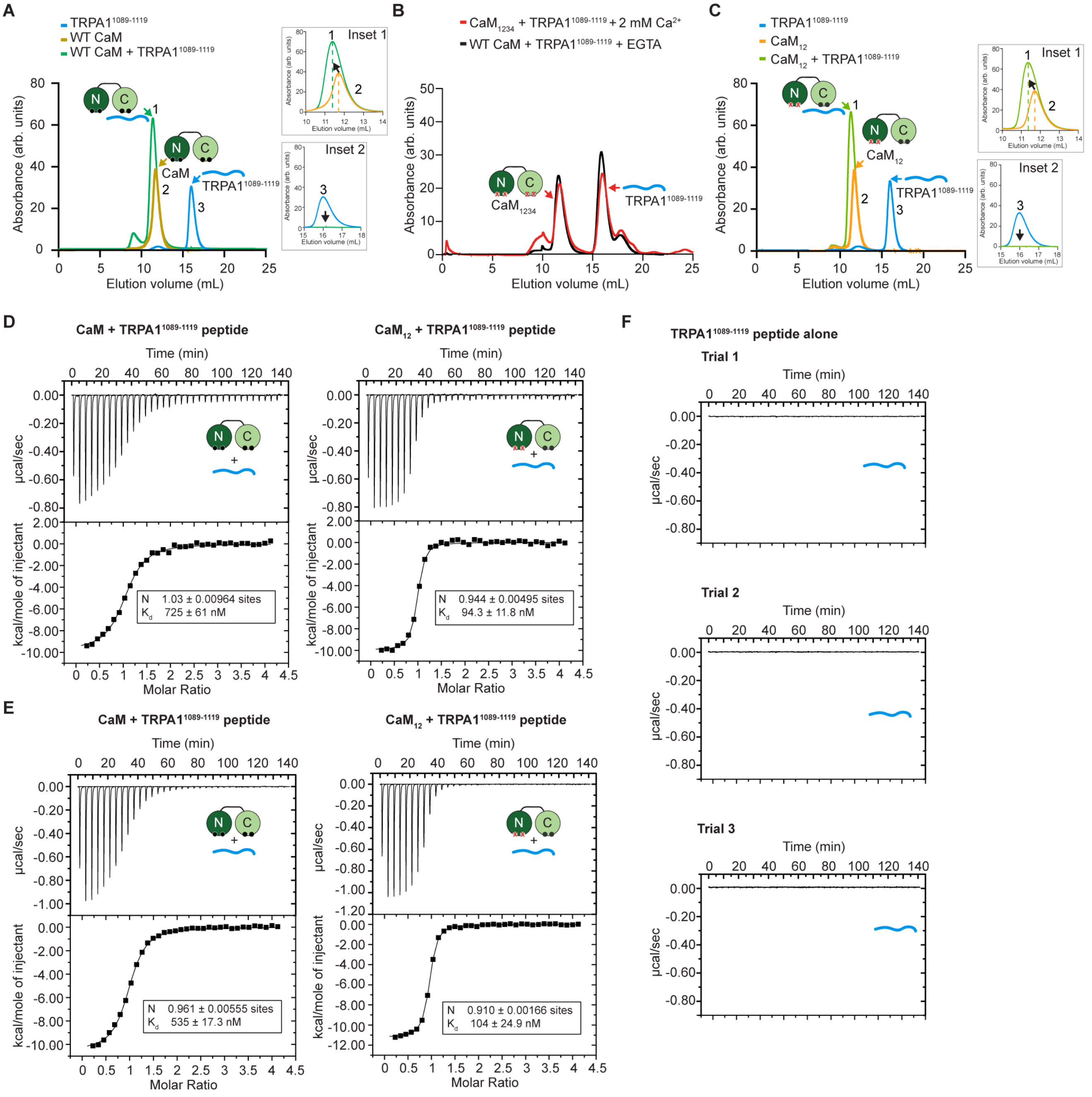


**Figure S5.** (**A**) Superdex 75 chromatograms of WT CaM alone (yellow), hTRPA1$^{1089\text{-}1119}$ peptide alone (blue), or WT CaM with hTRPA1$^{1089\text{-}1119}$ in the presence of 2 mM $Ca^{2+}$ (green). (**B**) Superdex 75 chromatogram of $CaM_{1234}$ with hTRPA1$^{1089\text{-}1119}$ in the presence of 2 mM $Ca^{2+}$ (red) plotted over WT CaM with hTRPA1$^{1089\text{-}1119}$ in the presence of 5 mM EGTA (black) from Fig. 4B. (**C**) Superdex 75 chromatograms of $CaM_{12}$ alone (orange), hTRPA1$^{1089\text{-}1119}$ alone (blue), and $CaM_{12}$ with hTRPA1$^{1089\text{-}1119}$ in the presence of 2 mM $Ca^{2+}$ (green). (A and C) Insets 1 focus on the CaM peak shift (to the left and increased UV signal) when bound to the hTRPA1$^{1089\text{-}1119}$ peptide. Insets 2 focus on the hTRPA1$^{1089\text{-}1119}$ peptide peak, which disappears when it binds to WT CaM or $CaM_{12}$. Chromatograms were generated from 100 µM protein for each construct. Data are representative of three independent replicates. (**D** and **E**) Replicates 2 (D) and 3 (E) of isothermal titration calorimetry plots of WT CaM (left) or $CaM_{12}$ (right) titrated by hTRPA1$^{1089\text{-}1119}$ at 2 mM $Ca^{2+}$ and fitted using the one-site binding model. Values for the number of binding sites (N) and the binding constant $K_d$ are shown. (**F**) Three control isothermal titration calorimetry plots for buffer titrated by TRPA1$^{1089\text{-}1119}$ at 2 mM $Ca^{2+}$.

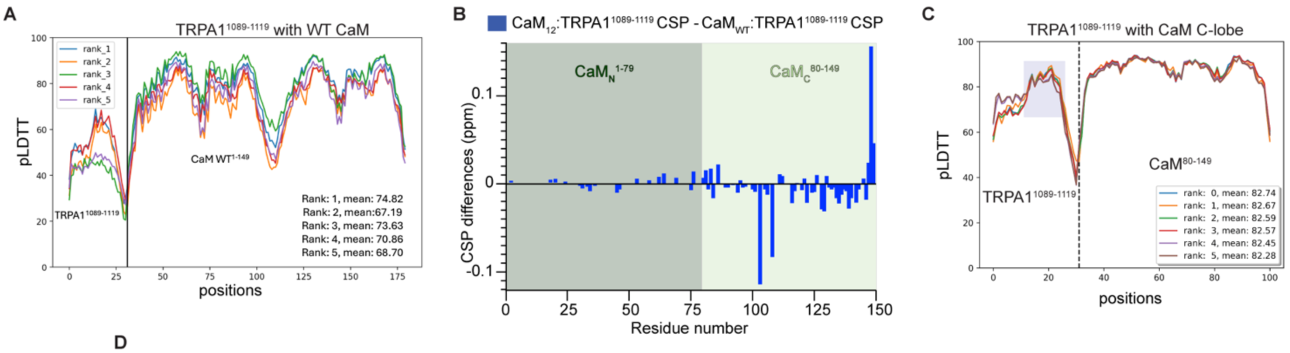


D

| PDB Accession Code | RMSD (Å) | Description of Structure | Calcification State | Lobe Engagement |
|---|---|---|---|---|
| 6YNU | 1.5 | CaM • CyaA translocation peptide | Fully | C-lobe only |
| 3G43 | 0.9 | CaM • Cav1.2 regulatory dimer | Fully | Lobes bind separate effectors |
| 3SUI | 1 | CaM • TRPV1 C-terminal peptide | Fully | Lobes engage same effector |
| 2W73 | 1.3 | CaM • Calcineurin A | Fully | Lobes bind separate effectors |
| 1IQ5 | 0.9 | CaM • CaMKK peptide | Fully | Lobes engage same effector |
| 1YR5 | 1 | CaM • DAP kinase peptide | Fully | Lobes engage same effector |
| 2VAY | 1.1 | CaM • Cav1.1 IQ peptide | Fully | Lobes engage same effector |
| 4M1L | 1 | CaM • IQCG IQ domain | Fully | Lobes engage same effector |
| 4DJC | 1.5 | CaM • Nav1.5 DIII-DIV linker peptide | Fully | C-lobe only |
| 2BE6 | 0.6 | CaM • CaV1.2 IQ domain | Fully | Lobes engage same effector |
| 7KL5 | 0.7 | CaM • Ryanodine receptor peptide | Fully | Lobes engage same effector |
| 3GOF | 1 | CaM • Macrophage nitric oxide synthase peptide | Fully | Lobes engage same effector |
| 5NIN | 1.6 | CaM • AKAP79 peptide | Fully | Lobes engage same effector |
| 3SJQ | 1.6 | CaM • SK channel CaMBD2 | Fully | Lobes bind separate effectors |
| 4EHQ | 1.7 | CaM • Orai1 CaM binding domain | Fully | C-lobe only |
| 5HIT | 1.4 | CaM • EAG1 channel C-terminal peptide | C-lobe only | C-lobe only |
| 3DVK | 1.5 | CaM • Cav2.3 IQ domain | Fully | Lobes engage same effector |
| 6SZ5 | 1.8 | CaM • NADPH oxidase 5 peptide | Fully | Lobes bind separate effectors |
| 4AQR | 1 | CaM • PMCA regulatory domain peptide | Fully | Lobes engage same effector |
| 6MUE | 1.8 | CaM • Nav1.4 IQ domain | Fully | C-lobe only |

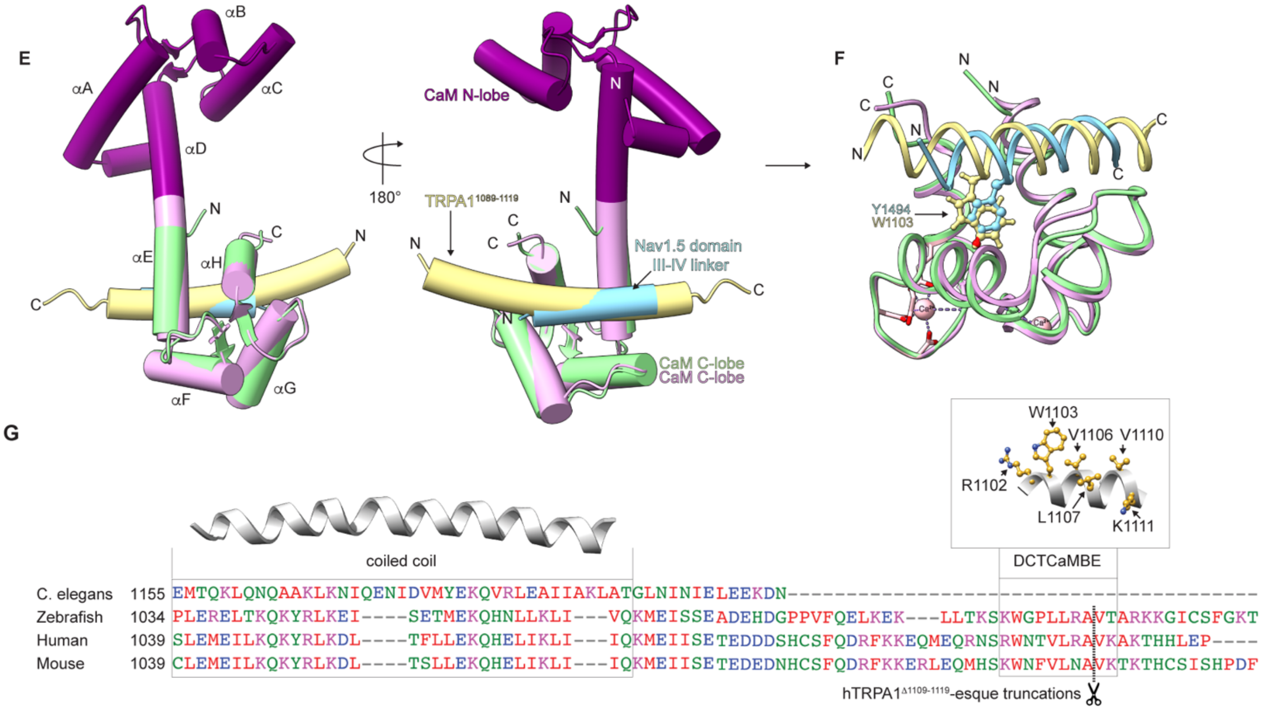


**Figure S6**. Structural models, analysis of the TRPA1 C-terminal tail CaM binding mode, and sequence conservation of the TRPA1 DCTCaMBE. (**A** and **C**) Confidence measurements (pLDDT plots) of AlphaFold2 Multimer models for the hTRPA1$^{1089\text{-}1119}$ peptide docked to WT hCaM (A) or the CaM C-lobe (C). For both, the top five generated models are plotted. Rank 1 (A) or 0 (C) models were used for models in Fig. 5A, 5E, 6A-C, and panels E and F. (**A**) hTRPA1$^{1089\text{-}1119}$ docked to the WT hCaM used in Fig. 5A. (**B**) CSP differences of the 1:1 $CaM_{WT}$:TRPA1$^{1089\text{-}1119}$ complex and the 1:1 $CaM_{12}$:TRPA1$^{1089\text{-}1119}$ complex from Fig. 5C as a function of residue number. Dark and light green shadings denote CaM N- and C-lobe residues, respectively. (**C**) hTRPA1$^{1089-1119}$ docked to the CaM C-lobe (residues 80-149) used in Fig. 5E, 6A-C, and panels E and F. (**D**) RMSD of hTRPA1$^{1089-1119}$ docked to the CaM C-lobe (residues 80-149) against all available structures. Calculations were performed by the Dali server using hTRPA1$^{1089-1119}$ docked to the CaM C-lobe as the search model in a heuristic PDB search. The top twenty results are listed with the structure description, lobe calcification state, and lobe engagement modes. (**E**) Ribbon diagram of hTRPA1$^{1089-1119}$ (yellow) docked to the CaM C-lobe (green) overlaid with the $Ca^{2+}$/CaM•Nav1.5 DIII-DIV linker peptide complex (CaM N- and C-lobes are indicated with dark and light purple, respectively, and the Nav1.5 peptide is indicated in blue). Models built with the rank 0 model from panel C or the CaM:Nav1.5 DIII-DIV linker peptide crystal structure (PDB: 4DJC) in ChimeraX. Models were aligned in real space. Two views are shown. (**F**) Ribbon diagram of CaM C-lobe:effector alignments from E with bulky hydrophobic residues that mediate interactions with the CaM C-lobe hydrophobic groove indicated for the Nav1.5 DIII-DIV linker peptide (blue, Y1494) or TRPA1$^{1089-1119}$ (yellow, W1103). $Ca^{2+}$ ions are indicated in pink. (**G**) Alignment of mouse, zebrafish TRPA1a isoform, and *C. elegans* TRPA1 with the hTRPA1 protein sequence for the coiled coil and structurally unresolved distal C-terminus. Regions of the coiled coil and the novel DCTCaMBE identified in this study are boxed and indicated above. Residues that contribute to CaM binding are indicated as balls and sticks in yellow. Alignment was built with Clustal Omega. The coiled coil helix was built with the human TRPA1 Cryo-EM structure (PDB: 6V9W) and the DCTCaMBE was built with the human TRPA1 AlphaFold deposited structure in ChimeraX. Scissors denote the hTRPA1$^{\Delta 1109-1119}$ truncation point.

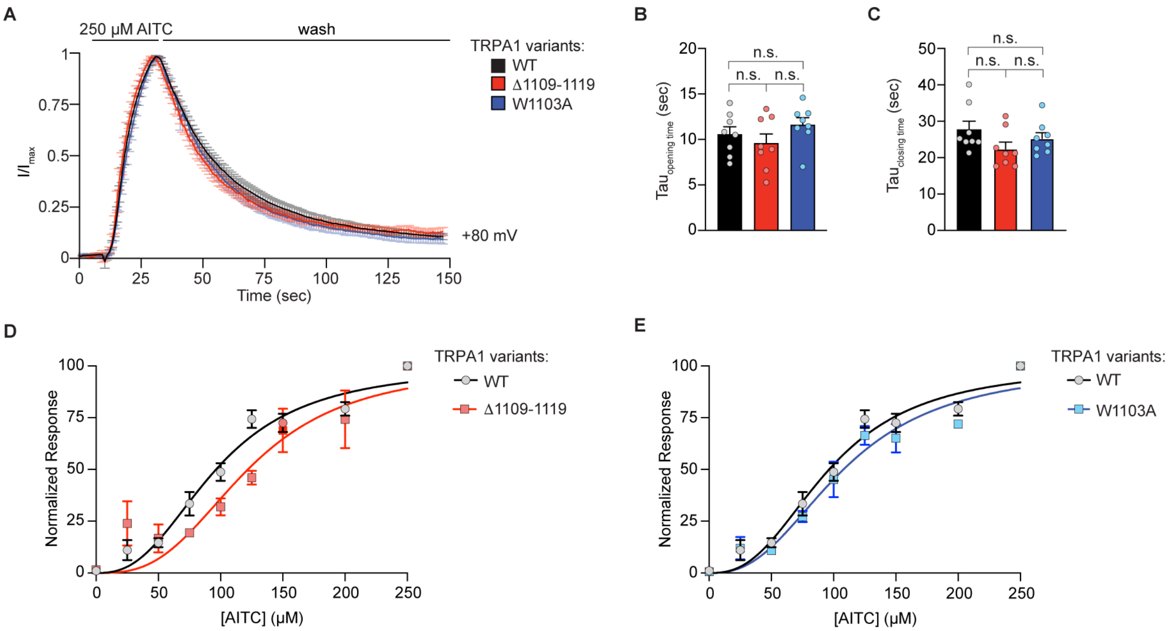


**Figure S7.** $Ca^{2+}$-free functional characterization of $Ca^{2+}$/CaM binding deficient TRPA1 mutants. (**A**) Average time-traces at +80 mV holding potential from oocytes expressing WT (black), Δ1109-1119 (red), or W1103A hTRPA1 (blue). n=8 independent oocytes. Error bars represent SEM across individual cell measurements. Current evoked with 250 μM AITC and then washed out. Extracellular solution contained no calcium. Currents (**I**) normalized to the maximum current ($I_{max}$) within each individual cell. (**B** and **C**) Calculated time constants of activation (B) and channel closing (C) at +80 mV from fitting data as in A to a single-exponential function. Data represent mean ± SEM. n.s. not significant p>0.05. n=8 oocytes per condition, one-way ANOVA with Tukey's *post hoc* analysis. (**D** and **E**) Dose-response curves of AITC-evoked responses from oocytes expressing WT (black, D and E), Δ1109-1119 (red, D), or W1103A hTRPA1 (blue, E). Evoked currents were normalized to maximum currents evoked by 250 μM AITC. Traces represent average ± SEM of normalized AITC-evoked currents from 5-8 independent oocytes per agonist concentration. Data were fit to a non-linear regression. $EC_{50}$ (95% CI) values are 96.3 μM for WT hTRPA1 (95% CI, 91.9-100.7 μM), 121.2 μM for hTRPA1$^{\Delta 1109\text{-}1119}$ (95% CI, 112.3-131.3 μM), and 107.9 μM for W1103A hTRPA1 (95% CI, 102.5-113.4 μM).

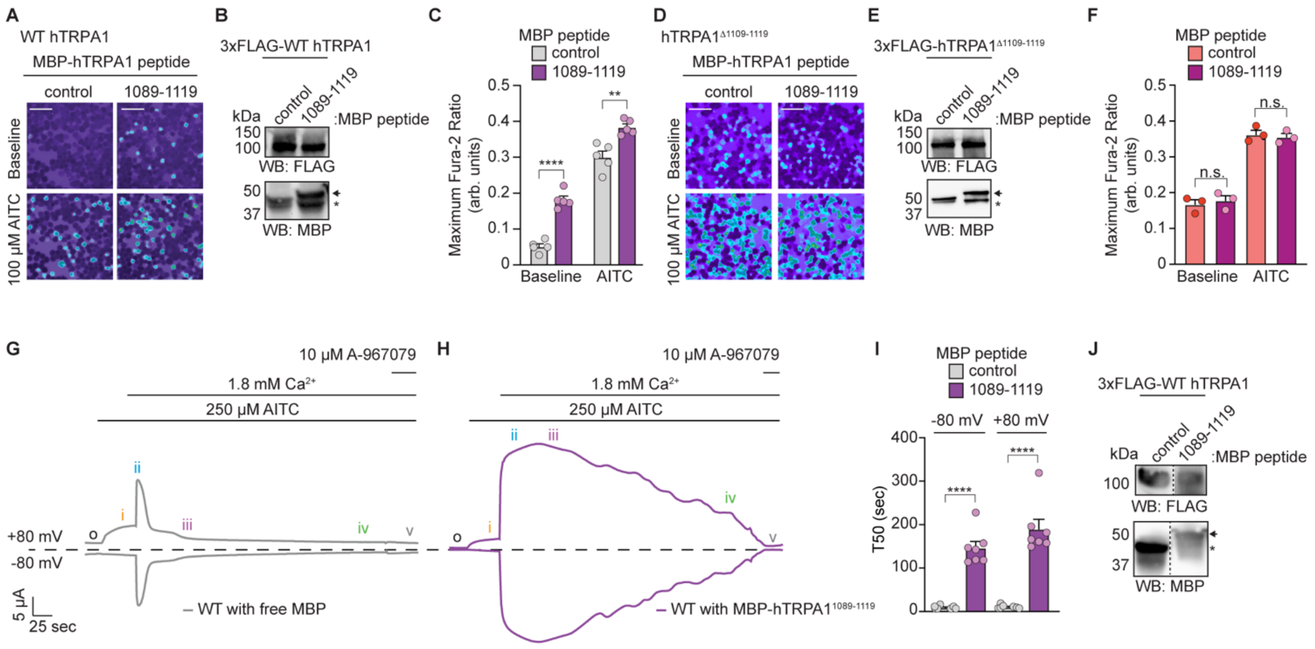


**Figure S8.** Endogenous CaM sequestration with a TRPA1 C-terminal peptide slows channel desensitization. (**A** and **D**) Ratiometric $Ca^{2+}$ imaging of HEK293T cells co-transfected with WT hTRPA1 (A) or hTRPA1$^{\Delta 1109\text{-}1119}$ (D) and free MBP (control) or MBP-hTRPA1$^{1089\text{-}1119}$. Cells were stimulated with 100 µM AITC. Images are representative of five (A) or three (D) independent experiments. Scale bars indicate 50 µm. (**B** and **E**) Representative immunoblotting analyses of the cells used for $Ca^{2+}$ imaging in A (B) or D (E). Samples were probed using an HRP-conjugated anti-FLAG antibody or an anti-MBP primary antibody and an HRP-conjugated anti-mouse secondary antibody. Asterisk (*) denotes free MBP. Arrow denotes MBP-tagged peptide. (**C** and **F**) Quantification of 100 µM AITC-evoked change in Fura-2 ratio of data from panel A (C) or D (F). (C) WT hTRPA1 co-expressed with free MBP indicated in grey, WT hTRPA1 co-expressed with MBP-hTRPA1$^{1089\text{-}1119}$ peptide indicated in purple. (F) hTRPA1$^{\Delta 1109\text{-}1119}$ co-expressed with free MBP indicated in salmon, hTRPA1$^{\Delta 1109\text{-}1119}$ co-expressed with MBP-hTRPA1$^{1089\text{-}1119}$ peptide indicated in plum. Data represent mean ± SEM. ****$p<0.0001$, **$p=0.0018$, n.s. not significant ($p>0.05$). n = 5 (C) or 3 (F) independent experiments, n ≥ 90 cells per transfection condition per experiment, two-tailed Student's t-test. (**G**-**H**) Representative time-traces at -80 and +80 mV holding potentials from oocytes expressing WT hTRPA1 and injected with 100 µM free MBP (G, grey) or MBP-hTRPA1$^{1089\text{-}1119}$ peptide (H, purple) one hour prior to recordings. Current evoked with 250 µM AITC in the absence (orange i) and presence (blue ii) of 1.8 mM extracellular $Ca^{2+}$. Channels were blocked with 10 µM A-967079 (grey v). Dashed line denotes 0 µA current. Protocol of condition application indicated above. (**I**) Calculated time constants of desensitization (T50) at -80 mV (left) and +80 mV (right) from fitting data as in G and H to a linear regression function. Data represent mean ± SEM. ****$p<0.0001$, two-tailed Student's t-test. n = 7 oocytes per injection type. Colors as in (C). (**J**) Western blot of lysates from oocytes used for recordings in G and H probed as in (B).
